# MAVE-NN: learning genotype-phenotype maps from multiplex assays of variant effect

**DOI:** 10.1101/2020.07.14.201475

**Authors:** Ammar Tareen, Mahdi Kooshkbaghi, Anna Posfai, William T. Ireland, David M. McCandlish, Justin B. Kinney

## Abstract

Multiplex assays of variant effect (MAVEs) are a family of methods that includes deep mutational scanning (DMS) experiments on proteins and massively parallel reporter assays (MPRAs) on gene regulatory sequences. However, a general strategy for inferring quantitative models of genotype-phenotype (G-P) maps from MAVE data is lacking. Here we introduce MAVE-NN, a neural-network-based Python package that implements a broadly applicable information-theoretic framework for learning G-P maps—including biophysically interpretable models—from MAVE datasets. We demonstrate MAVE-NN in multiple biological contexts, and highlight the ability of our approach to deconvolve mutational effects from otherwise confounding experimental nonlinearities and noise.

## Introduction

Over the last decade, the ability to quantitatively study genotype-phenotype (G-P) maps has been revolutionized by the development of multiplex assays of variant effect (MAVEs), which can measure molecular phenotypes for thousands to millions of genotypic variants in parallel.^1,2^ MAVE is an umbrella term that describes a diverse set of experimental methods, three examples of which are illustrated in **Fig. 1**. Deep mutational scanning (DMS) experiments^3^ are a type of MAVE commonly used to study protein sequence-function relationships. These assays work by linking variant proteins to their coding sequences, either directly or indirectly, then using deep sequencing to assay which variants survive a process of activity-dependent selection (e.g., **Fig. 1a**). Massively parallel reporter assays (MPRAs) are another major class of MAVE, and are commonly used to study DNA or RNA sequences that regulate gene expression at a variety of steps, including transcription, mRNA splicing, cleavage and polyadenylation, translation, and mRNA decay.^4–7^ MPRAs typically rely on either an RNA-seq readout of barcode abundances (**Fig. 1c**) or the sorting of cells expressing a fluorescent reporter gene (**Fig. 1e**).

**Figure 1.**
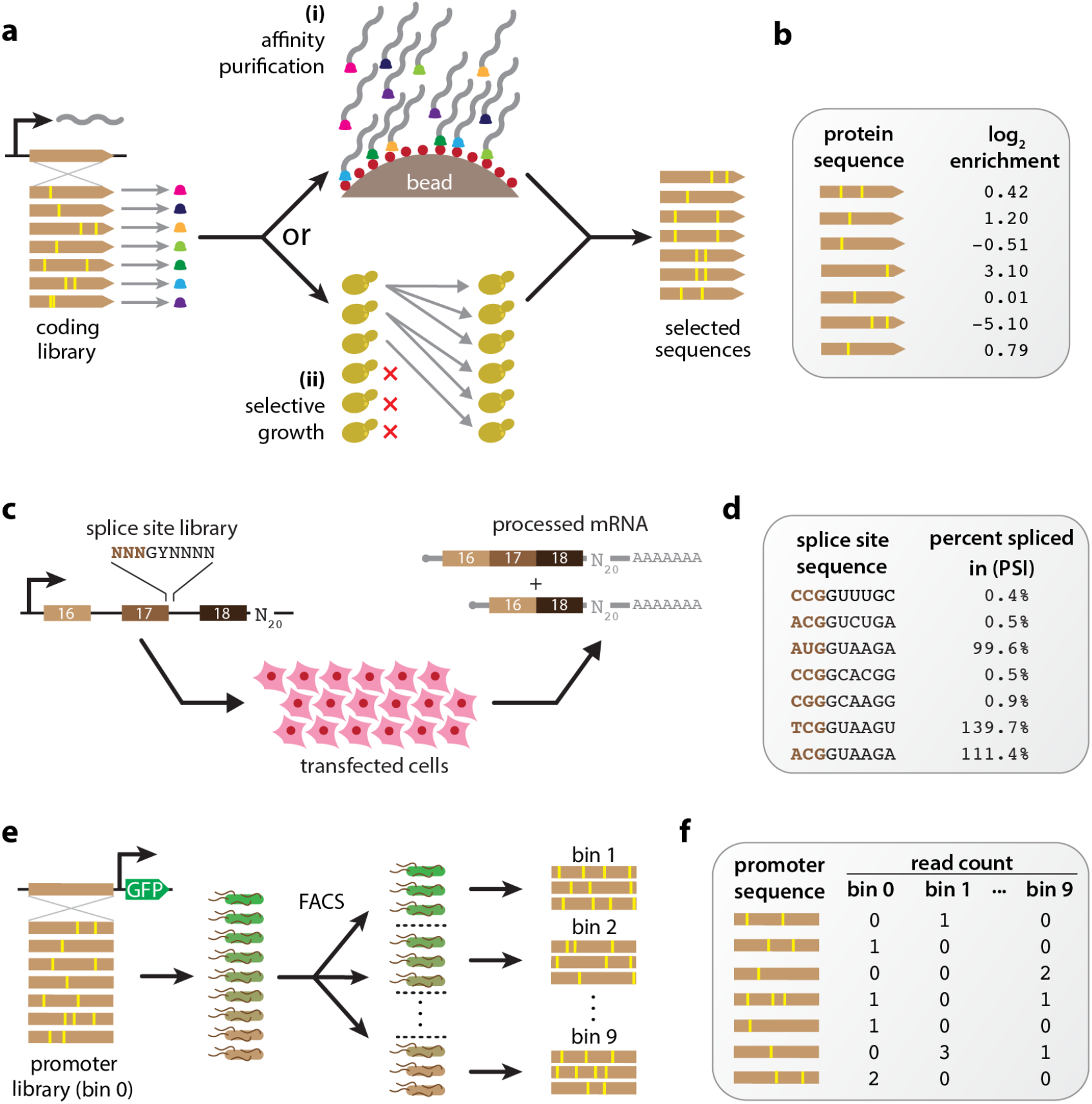
Examples illustrating the diversity of MAVEs. (**a**) DMS assays using either affinity purification or selective growth. **(i)** The DMS assay of Olson et al.^35^ used a library of variant GB1 proteins covalently linked to their coding mRNAs via mRNA display. Functional GB1 proteins were then enriched using IgG beads. **(ii)** The DMS studies of Seuma et al.^36^ and Bolognesi et al.^37^ used selective growth in genetically modified *Saccharomyces cerevisiae* cells to respectively assay the functionality of variant Aβ and TDP-43 proteins. In all three experiments, deep sequencing was used to determine an enrichment ratio for each protein variant. (**b**) The resulting DMS dataset consists of variant protein sequences and their corresponding log enrichment values. (**c**) The MPSA of Wong et al..^38^ A library of 3-exon minigenes was constructed from exons 16, 17, and 18 of *BRCA2*, with each minigene having a variant 5’ss at exon 17 and a random 20 nt barcode in the 3’ UTR. This library was transfected into HeLa cells, and deep sequencing was used to quantify mRNA isoform abundance. (**d**) The resulting MPSA dataset comprises variant 5’ss with (noisy) PSI values. (**e**) The sort-seq MPRA of Kinney et al..^16^ A plasmid library was generated in which randomly mutagenized versions of the *Escherichia coli lac* promoter drove the expression of GFP. Cells carrying these plasmids were sorted using FACS, and the variant promoters in each bin of sorted cells as well as the initial library were sequenced. (**f**) The resulting dataset comprises a list of variant promoter sequences, as well as a matrix of counts for each variant in each FACS bin. MAVE: multiplex assay of variant effect; DMS: deep mutational scanning; MPSA: massively parallel splicing assay; 5’ss: 5’ splice site(s); PSI: percent spliced in; GFP: green fluorescent protein; FACS: fluorescence-activated cell sorting.

Most computational methods for analyzing MAVE data have focused on accurately quantifying the activity of individual assayed sequences.^8–14^ However, MAVE measurements like enrichment ratios or cellular fluorescence levels usually cannot be interpreted as providing direct quantification of biologically meaningful activities, due to the presence of experiment-specific nonlinearities and noise. Moreover, MAVE data is usually incomplete, as one often wishes to understand G-P maps over vastly larger regions of sequence space than can be exhaustively assayed. The explicit quantitative modeling of G-P maps can address both the indirectness and incompleteness of MAVE measurements.^1,15^ The goal here is to determine a mathematical function that, given a sequence as input, will return a quantitative value for that sequence’s molecular phenotype. Such quantitative modeling has been of great interest since the earliest MAVE methods were developed,^16–18^ but no general-use software has yet been described for inferring G-P maps of arbitrary functional form from MAVE data.

Here we introduce a unified conceptual framework for the quantitative modeling of MAVE data. This framework is based on the use of latent phenotype models, which assume that each assayed sequence has a well-defined latent phenotype (specified by the G-P map), of which the MAVE experiment provides an indirect readout (described by the measurement process). The quantitative forms of both the G-P map and the measurement process are then inferred from MAVE data simultaneously. We further introduce an information-theoretic approach for separately assessing the performance of the G-P map and the measurement process components of latent phenotype models. This strategy is implemented in an easy-to-use open-source Python package called MAVE-NN, which represents latent phenotype models as neural networks and infers the parameters of these models from MAVE data using a TensorFlow 2 backend.^19^

In what follows, we expand on this unified MAVE modeling strategy and apply it to a diverse array of DMS and MPRA datasets. Doing so, we find that MAVE-NN provides substantial advantages over other MAVE modeling approaches. Our results also highlight the substantial benefits of including sequence variants with multiple mutations in assayed sequence libraries, as doing so allows MAVE-NN to deconvolve the features of the G-P map from potentially confounding effects of experimental nonlinearities and noise. Importantly, we find that including just a modest number of multiple-mutation variants in a MAVE experiment can be beneficial even when one is primarily interested in the effects of single mutations. Finally, we illustrate how the ability of MAVE-NN to train custom G-P maps can shed light on the biophysical mechanisms of gene regulation.

## Results

### Latent phenotype modeling strategy

MAVE-NN supports the analysis of MAVE data on DNA, RNA, and protein sequences, and can accommodate either continuous or discrete measurement values. Given a set of sequence-measurement pairs, MAVE-NN aims to infer a probabilistic mapping from sequence to measurement. Our primary enabling assumption, which is encoded in the structure of the latent phenotype model (**Fig. 2a**), is that this mapping occurs in two stages. Each sequence is first mapped to a latent phenotype by a deterministic G-P map, then this latent phenotype is mapped to possible measurement values via a stochastic measurement process. During training, the G-P map and measurement process are simultaneously learned by maximizing a regularized form of likelihood.

**Figure 2.**
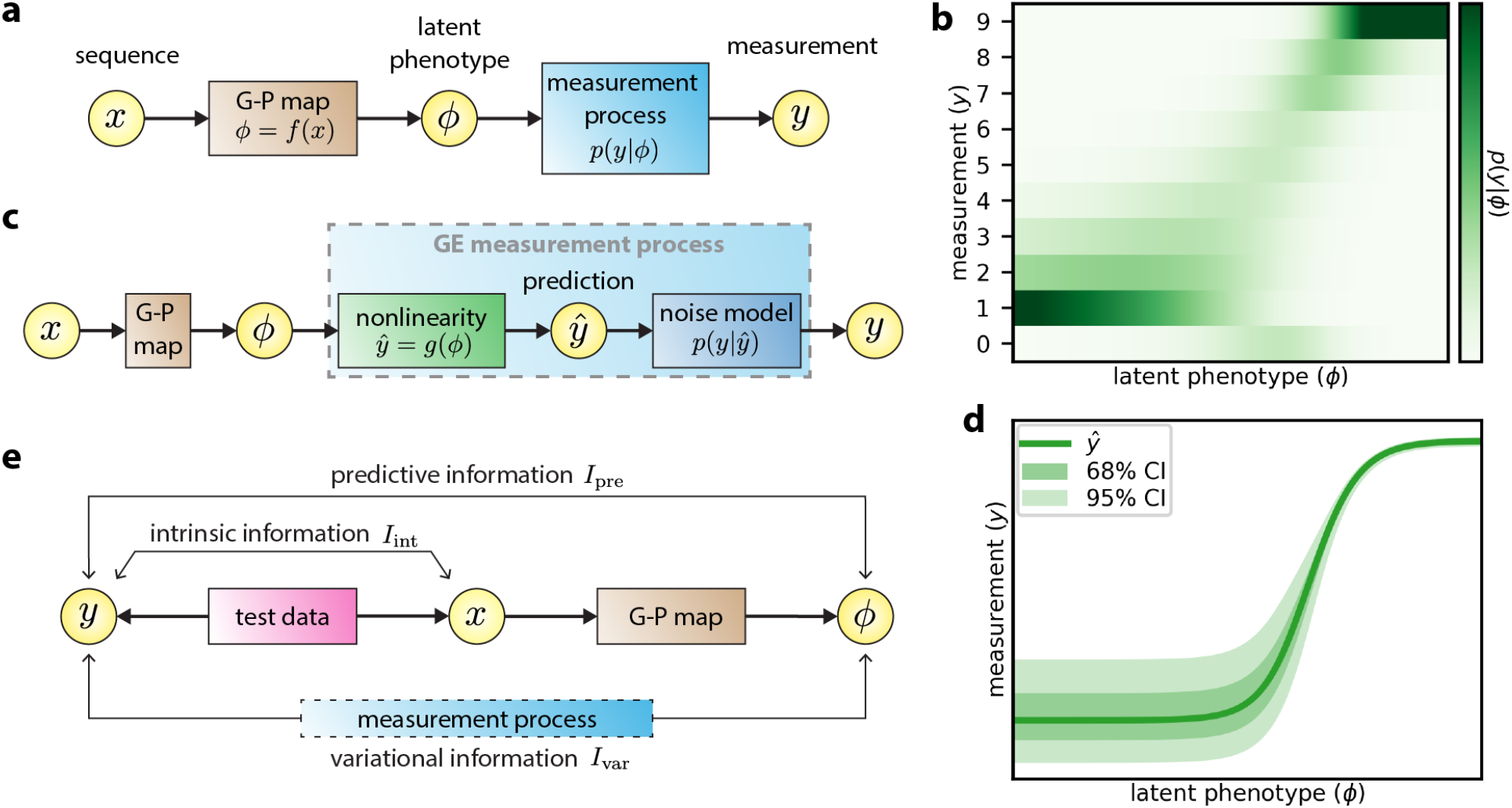
MAVE-NN quantitative modeling strategy. (**a**) Structure of latent phenotype models. A G-P map *f*(*x*) maps each sequence *x* to a latent phenotype *ϕ*, after which a measurement process *p*(*y*|*ϕ*) determines the measurement *y*. (**b**) Example of an MPA measurement process inferred from the sort-seq MPRA data of Kinney et al..^16^ MPA measurement processes are used when *y* values are discrete. (**c**) Structure of a GE regression model, which is used when *y* is continuous. A GE measurement process assumes that the mode of *p*(*y*|*ϕ*), called the prediction *ŷ*, is given by a nonlinear function *g*(*ϕ*), and the scatter about this mode is described by a noise model *p*(*y*|*ŷ*). (**d**) Example of a GE measurement process inferred from the DMS data of Olson et al..^35^ Shown are the nonlinearity, the 68% CI, and the 95% CI. (**e**) Information-theoretic quantities used to assess model performance. Intrinsic information, *I*_int_, is the mutual information between sequences *x* and measurements *y*. Predictive information, *I*_pre_, is the mutual information between measurements *y* and the latent phenotype values *ϕ* assigned by a model. Variational information, *I*_var_, is a linear transformation of log likelihood. The inequality *I*_int_ ≥ *I*_pre_ ≥ *I*_var_ always holds on test data (modulo finite data uncertainties), with *I*_int_ = *I*_pre_ when the G-P map is correct, and *I*_pre_ = *I*_var_ when the measurement process correctly describes the distribution of *y* conditioned on *ϕ*. G-P: genotype-phenotype; MPA: measurement process agnostic; GE: global epistasis; CI: confidence interval.

MAVE-NN includes four types of built-in G-P maps: additive, neighbor, pairwise, and black box. Additive G-P maps assume that each character at each position within a sequence contributes independently to the latent phenotype. Neighbor G-P maps incorporate interactions between adjacent (i.e., nearest-neighbor) characters in a sequence, while pairwise G-P maps include interactions between all pairs of characters in a sequence regardless of the distance separating the characters in each pair. Black box G-P maps have the form of a densely connected multilayer perceptron, the specific architecture of which can be controlled by the user. MAVE-NN also supports custom G-P maps that can be used, e.g., to represent specific biophysical hypotheses about the mechanisms of sequence function.

To handle both discrete and continuous measurement values, two different strategies for modeling measurement processes are provided. Measurement process agnostic (MPA) regression uses techniques from the biophysics literature^15,16,20,21^ to analyze MAVE datasets that report discrete measurements. Here the measurement process is represented by an overparameterized neural network that takes the latent phenotype value as input and outputs the probability of each possible measurement value (**Fig. 2b**). Global epistasis (GE) regression (**Fig. 2c**), by contrast, leverages ideas previously developed in the evolution literature for analyzing datasets that contain continuous measurements,^22–25^ and is becoming an increasingly popular strategy for modeling DMS data.^26–28^ Here, the latent phenotype is nonlinearly mapped to a prediction that represents the most probable measurement value. A noise model is then used to describe the distribution of likely deviations from this prediction. MAVE-NN supports both homoscedastic and heteroscedastic noise models based on three different classes of probability distribution: Gaussian, Cauchy, and skewed-t. We note that the skewed-t distribution, introduced by Jones and Faddy,^29^ reduces to Gaussian and Cauchy distributions in certain limits while also accommodating asymmetric experimental noise. **Fig. 2d** shows an example of a GE measurement process with a heteroscedastic skewed-t noise model.

Readers should note that the current implementation of MAVE-NN places certain constraints on input data and model architecture. Input sequences must be the same length, and when analyzing continuous data, only scalar measurements (as opposed to vectors of multiple measurements) can be used to train models. In addition, because our method for learning the form of experimental nonlinearities depends on observing how multiple mutations combine, MAVE-NN’s functionality is more limited when analyzing MAVE libraries that comprise only single-mutation variants. More information on these constraints and the reasons behind them can be found below in the section “Constraints on datasets and models”.

### Information-theoretic measures of model performance

We further propose three distinct quantities for assessing the performance of latent phenotype models: intrinsic information, predictive information, and variational information (**Fig. 2e**). These quantities come from information theory and are motivated by thinking of G-P maps in terms of information compression. In information theory, a quantity called mutual information quantifies the amount of information that the value of one variable communicates about the value of another.^30,31^ Mutual information is symmetric, nonnegative, is measured in units of “bits”, and is equal to 0 bits only if the two variables are independent. Alternatively, if knowing the value of one variable allows you to narrow down the value of the other variable to one of two possibilities that would otherwise be equally likely, the mutual information between these two variables will be 1.0 bits. If the value is narrowed down to one of four otherwise equally likely possibilities, the mutual information will be 2.0 bits. Narrowing down to one of with eight possibilities will yield 3.0 bits, and so on. But importantly, mutual information does not require that the relationship between two variables in question be so clean cut, and mutual information can in fact be computed between any two types of variables—discrete, continuous, multi-dimensional, etc.. This property makes the information-based quantities we propose applicable to all MAVE datasets, regardless of the specific type of experimental readout used. By contrast, many of the standard model performance metrics have restricted domains of applicability: accuracy can only be applied to data with discrete labels, *R*^2^ can only be applied to data with univariate continuous labels, and so on. We note, however, that estimating mutual information and related quantities from finite data is nontrivial and that MAVE-NN uses a variety of approaches to do this.

Intrinsic information, *I*_int_, is the mutual information between the sequences and measurements contained within a MAVE dataset. This quantity provides a benchmark against which to compare the performance of inferred G-P maps. Predictive information, *I*_pre_, is the mutual information between MAVE measurements and the latent phenotype values predicted by a G-P map of interest. This quantifies how well the G-P map preserves sequence-encoded information that is determinative of experimental measurements. When evaluated on test data, *I*_pre_ is bounded above by *I*_int_, and equality is realized only when the latent phenotype losslessly encodes relevant sequence-encoded information. Variational information, *I*_var_, is a linear transformation of log likelihood that provides a variational lower bound on *I*_pre_.^32–34^ The difference between *I*_pre_ and *I*_var_ quantifies how accurately the inferred measurement process matches the observed distribution of measurements and latent phenotypes (see **Supplemental Information**).

MAVE-NN infers model parameters by maximizing a (lightly) regularized form of likelihood. These computations are performed using the standard backpropagation-based training algorithms provided within the TensorFlow 2 backend. With certain caveats noted (see **Methods**), this optimization procedure maximizes *I*_pre_ while avoiding the costly estimates of mutual information at each iteration that have hindered the adoption of previous mutual-information-based modeling strategies.^16^

### Application: deep mutational scanning assays

We now demonstrate the capabilities of MAVE-NN on three DMS datasets, starting with the study of Olson et al.^35^ on pairwise epistasis in protein G. Here the authors measured the effects of all single and nearly all double mutations to residues 2-56 of the IgG binding domain. This domain, called GB1, has long served as a model system for studying protein sequence-function relationships. To assay the binding of GB1 variants to IgG, the authors combined mRNA display with ultra-high-throughput DNA sequencing (**Fig. 1a**). The resulting dataset reports log enrichment values for all 1,045 single- and 530,737 double-mutant GB1 variants (**Fig. 1b**).

Inspired by the work of Otwinowski et al.,^25^ we used MAVE-NN to infer a latent phenotype model comprising an additive G-P map and a GE measurement process. This inference procedure required only about 5 minutes on a single node of a computer cluster (**Supplemental Fig. S1**). **Fig. 3a** illustrates the inferred additive G-P map via the effects that every possible single-residue mutation has on the latent phenotype. From this heatmap of additive effects, we can immediately identify all of the critical GB1 residues, including the IgG interacting residues at 27, 31, and 43.^35^ We also observe that missense mutations to proline throughout the GB1 domain tend to negatively impact IgG binding, as expected due to this amino acid’s exceptional conformational rigidity. **Fig. 3b** illustrates the corresponding GE measurement process, revealing a sigmoidal relationship between log enrichment measurements and the latent phenotype values predicted by the G-P map. Nonlinearities like this are ubiquitous in DMS data due to the presence of background and saturation effects. Unless they are explicitly accounted for in one’s quantitative modeling efforts, as they are here, these nonlinearities can greatly distort the parameters of inferred G-P maps. **Fig. 3c** shows that accounting for this nonlinearity yields predictions that correlate quite well with measurement values. Moreover, every latent phenotype model inferred by MAVE-NN can be used as a MAVE dataset simulator (see **Methods**). By analyzing simulated data generated by our inferred model for this GB1 experiment, we further observed that MAVE-NN can accurately and robustly recover the GE nonlinearity and ground-truth G-P map parameters (**Supplementary Fig. S1**).

**Figure 3.**
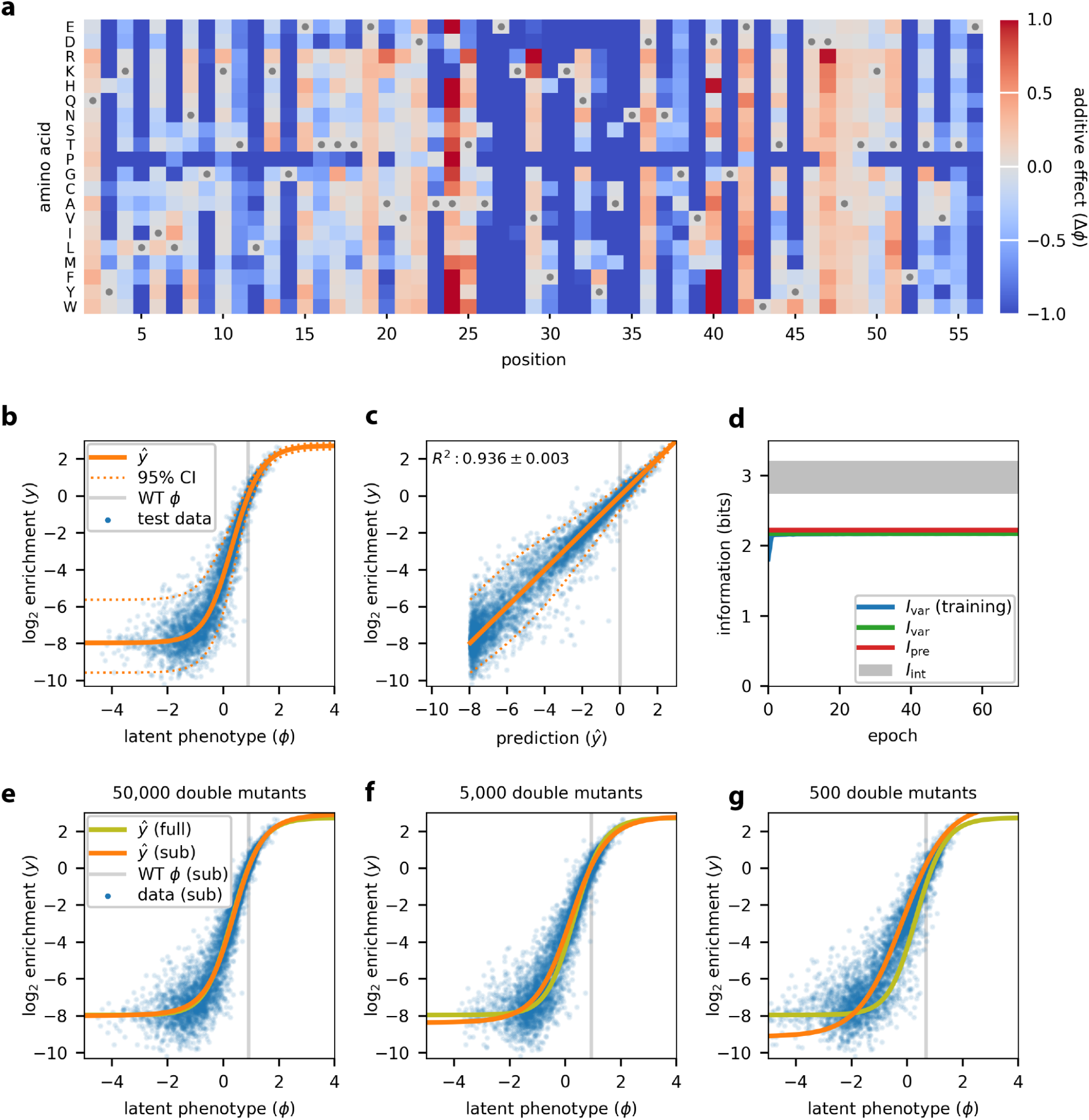
Analysis of DMS data for protein GB1. MAVE-NN was used to infer a latent phenotype model, consisting of an additive G-P map and a GE measurement process having a heteroskedastic skewed-t noise model, from the DMS data of Olson et al..^35^ All 530,737 pairwise variants reported for positions 2 to 56 of the GB1 domain were analyzed. Data were split 90:5:5 into training, validation, and test sets. (**a**) The G-P map parameters inferred from all pairwise variants. Gray dots indicate wildtype residues. Amino acids are ordered as in Olson et al..^35^ (**b**) GE plot showing measurements versus predicted latent phenotype values for 5,000 randomly selected test-set sequences (blue dots), alongside the inferred nonlinearity (solid orange line) and the 95% CI (dashed lines) of the noise model. Gray line indicates the latent phenotype value of the wildtype sequence. (**c**) Measurements plotted against *ŷ* predictions for these same sequences. Dashed lines indicate the 95% CI of the noise model. Gray line indicates the wildtype sequence *ŷ*. (**d**) Corresponding information metrics computed during model training (using training data) or for the final model (using test data); uncertainties in these estimates are roughly the width of the plotted lines. Gray shaded area indicates allowed values for intrinsic information based on upper and lower bounds estimated as described in **Methods**. (**e-g**) Test set predictions (blue dots) and GE nonlinearities (orange lines) for models trained using subsets of the GB1 data containing all single mutants and 50,000 (**e**), 5,000 (**f**), or 500 (**g**) double mutants. The GE nonlinearity from panel **b** is shown for reference (yellow-green lines). Uncertainties reflect standard errors. GE: global epistasis; G-P: genotype-phenotype; CI: confidence interval.

**Fig. 3d** summarizes the values of our information-theoretic metrics for model performance. On held-out test data, we find that *I*_var_ = 2.178 ± 0.027 bits and *I*_pre_ = 2.225 ± 0.017 bits. The similarity of these two values suggests that the inferred GE measurement process, which includes a heteroscedastic skewed-t noise model, has nearly sufficient accuracy to fully describe the distribution of residuals. We further find that 2.741 ± 0.013 bits ≤ *I*_int_ ≤ 3.215 ± 0.007 bits (see **Methods**), meaning that the inferred G-P map accounts for 69%-81% of the total sequence-dependent information in the dataset. While this performance is impressive, the additive G-P map evidently misses some relevant sequence features. This observation motivates the more complex biophysical model for GB1 discussed later in **Results**.

The ability of MAVE-NN to deconvolve experimental nonlinearities from additive G-P maps requires that some of the assayed sequences contain multiple mutations. This is because such nonlinearities are inferred by reconciling the effects of single mutations with the effects observed for combinations of two or more mutations. To investigate how many multiple-mutation variants are required, we performed GE inference on subsets of the GB1 dataset containing all 1,045 single-mutation sequences and either 50,000, 5,000, or 500 double-mutation sequences (see **Methods**). The shapes of the resulting GE nonlinearities are illustrated in **Figs. 3e-g**. Remarkably, MAVE-NN is able to recover the underlying nonlinearity using only about 500 randomly selected double mutants, which represent only ∼0.1% of all possible double mutants. The analysis of simulated data also supports the ability to accurately recover ground-truth model predictions using highly reduced datasets (**Supplemental Fig. S1**). These findings have important implications for the design of DMS experiments: even if one only wants to determine an additive G-P map, including a modest number of multiple-mutation sequences in the assayed library is often advisable because it may allow the removal of artifactual nonlinearities.

To test the capabilities of MAVE-NN on less complete DMS datasets, we analyzed recent experiments on amyloid beta (Aβ)^36^ and TDP-43,^37^ both of which exhibit aggregation behavior in the context of neurodegenerative diseases. In these experiments, protein functionality was assayed using selective growth in genetically modified *Saccaromyces cerevisiae*: Seuma et al.^36^ performed a selection against Aβ toxicity, whereas Bolognesi et al.^37^ positively selected for TDP-43 aggregation. Like with GB1, the variant libraries used in these two experiments included a substantial number of multiple-mutation sequences: 499 single- and 15,567 double-mutation sequences for Aβ; 1,266 single- and 56,730 double-mutation sequences for TDP-43. But unlike with GB1, these datasets are highly incomplete due to the use of mutagenic PCR (for Aβ) or doped oligo synthesis (TDP-43) for variant library construction.

We used MAVE-NN to infer additive G-P maps from these two datasets, adopting the same type of latent phenotype model used for GB1. **Fig. 4a** illustrates the additive G-P map inferred from aggregation measurements of Aβ variants. In agreement with the original study, we see that most amino acid mutations between positions 30-40 have a negative effect on nucleation, suggesting that this region plays a major role in nucleation behavior. **Fig. 4b** shows the corresponding measurement process (see also **Supplemental Information Fig. S2**). Even though these data are much sparser than the GB1 data, the inferred model performs well on held-out test data (*I*_var_ = 1.142 ± 0.065 bits, *I*_pre_ = 1.187 ± 0.050 bits, *R*^2^ = 0.763 ± 0.024). Similarly, **Figs. 4c-d** show the G-P map parameters and GE measurement process inferred from toxicity measurements of TDP-43 variants, revealing among other things the toxicity-determining hot-spot observed by Bolognesi et al.^37^ at positions 310-340. The resulting latent phenotype model performs well on held-out test data (*I*_var_ = 1.834 ± 0.035 bits, *I*_pre_ = 1.994 ± 0.023 bits, *R*^2^ = 0.914 ± 0.007).

**Figure 4.**
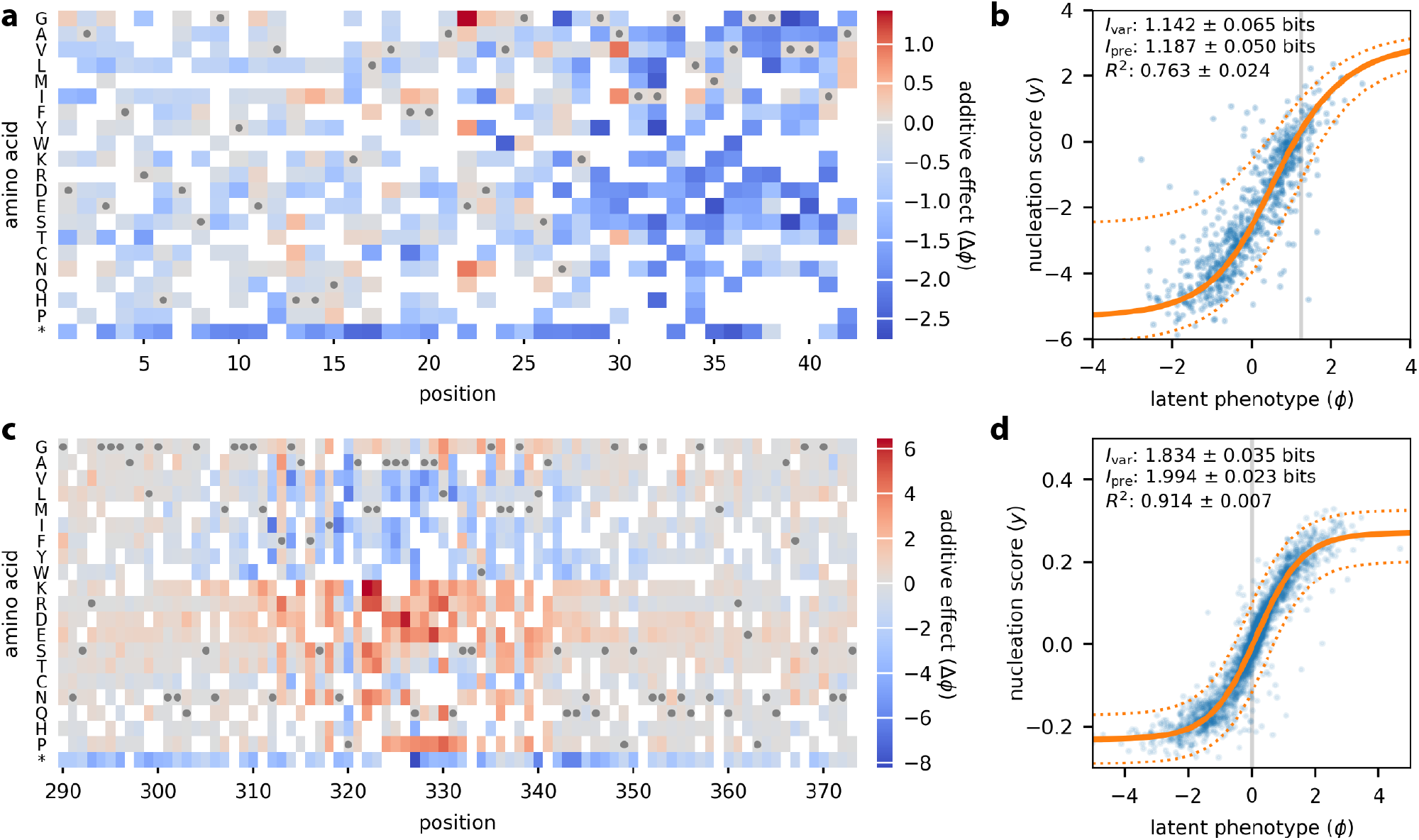
Analysis of DMS data for Aβ and TDP-43. (**a**,**b**) Seuma et al.^36^ measured nucleation scores for 499 single mutants and 15,567 double mutants of Aβ. These data were used to train a latent phenotype model comprising (**a**) an additive G-P map and (**b**) a GE measurement process with a heteroskedastic skewed-t noise model. (**c**,**d**) Bolognesi et al.^37^ measured toxicity scores for 1,266 single mutants and 56,730 double mutants of TDP-43. The resulting data were used to train (**c**) an additive G-P map and (**d**) a GE measurement process of the same form as in panel **b**. In both cases, data were split 90:5:5 into training, validation, and test sets. In (**a**,**c**), gray dots indicate the wildtype sequence, amino acids are ordered as in the original publications, and * indicates a stop codon. White squares [355/882 (40.2%) for Aβ; 433/1764 (24.5%) for TDP-43] indicate residues that were not observed in the training set and thus could not be assigned values for their additive effects. In (**b**,**d**), blue dots indicate latent phenotype values versus measurements for held-out test data, gray line indicates the latent phenotype value of the wildtype sequence, solid orange line indicates the GE nonlinearity, and dashed orange lines indicate a corresponding 95% CI for the inferred noise model. Values for *I*_var_, *I*_pre_, and *R*^2^ (between *y* and *ŷ*) are also shown. Uncertainties reflect standard errors. **Supplemental Fig. S2** shows measurements plotted against the *ŷ* predictions of these models. Aβ: amyloid beta; TDP-43: TAR DNA-binding protein 43; G-P: genotype-phenotype; GE: global epistasis; CI: confidence interval.

### Application: a massively parallel splicing assay

Exon/intron boundaries are defined by 5’ splice sites (5’ss), which bind the U1 snRNP during the initial stages of spliceosome assembly. To investigate how 5’ss sequence quantitatively controls alternative mRNA splicing, Wong et al.^38^ used a massively parallel splicing assay (MPSA) to measure percent-spliced-in (PSI) values for nearly all 32,768 possible 5’ss of the form NNN/GYNNNN in three different genetic contexts (**Fig. 1c**,**d**). Applying MAVE-NN to data from the BRCA2 exon 17 context, we inferred four different types of G-P maps: additive, neighbor, pairwise, and black box. As with GB1, these G-P maps were each inferred using GE regression with a heteroscedastic skewed-t noise model. For comparison, we also inferred an additive G-P map using the epistasis package of Sailer and Harms.^24^

**Fig. 5a** compares the performance of these G-P map models on held-out test data, while **Figs. 5b-d** illustrate the corresponding inferred measurement processes. We observe that the additive G-P map inferred using the epistasis package^24^ exhibits less predictive information (*I*_pre_ = 0.180 ± 0.011 bits) than the additive G-P map found using MAVE-NN (*P =*3.8 × 10^−6^, two-sided z-test). This is likely because the epistasis package estimates the parameters of the additive G-P map prior to estimating the GE nonlinearity. We also note that, while the epistasis package provides a variety of options for modeling the GE nonlinearity, none of these options appear to work as well as our mixture-of-sigmoids approach (compare **Figs. 5b**,**c**). This finding again demonstrates that the accurate inference of G-P maps requires the explicit and simultaneous modeling of experimental nonlinearities.

**Figure 5.**
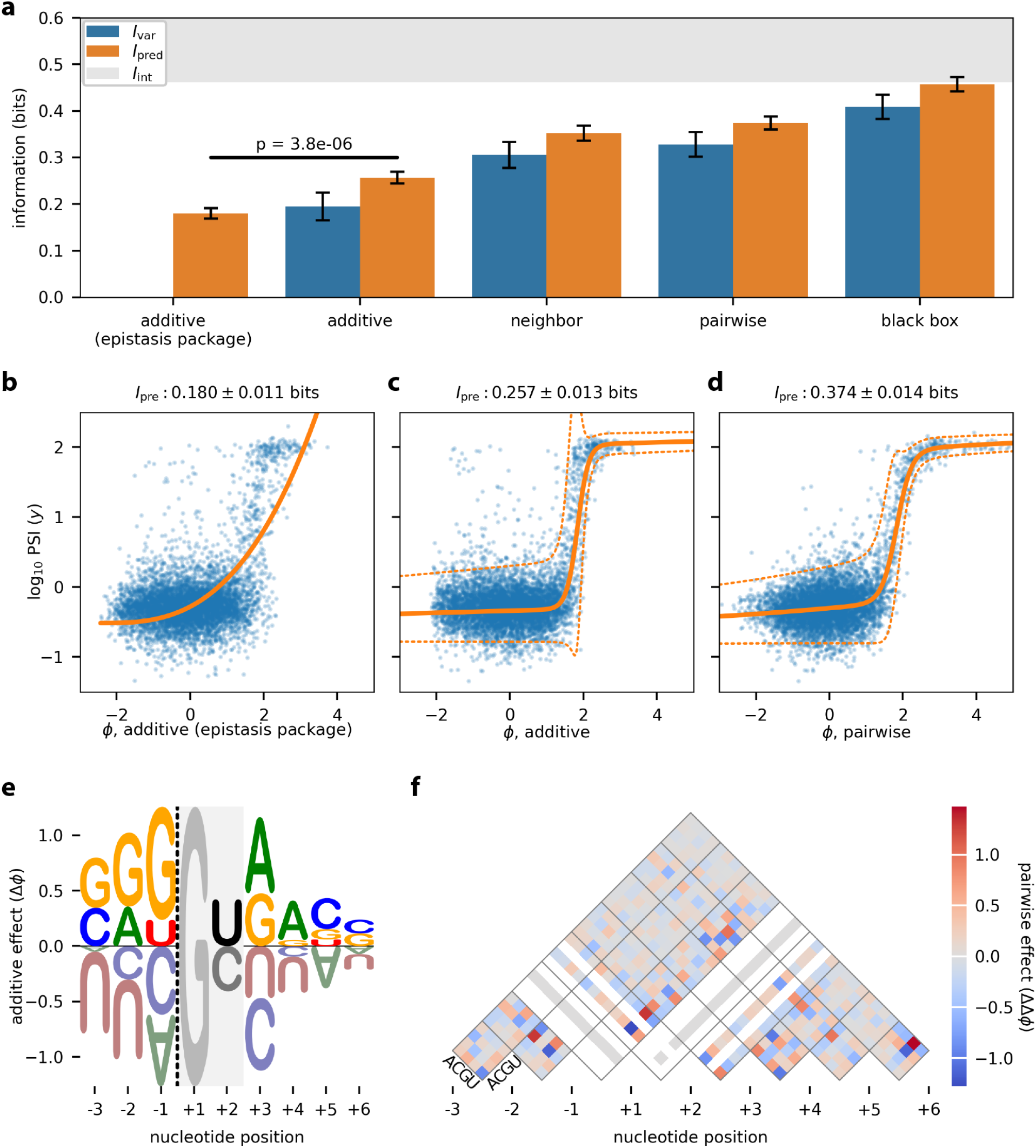
Analysis of MPSA data from Wong et al..^38^ This dataset reports PSI values, measured in the *BRCA2* exon 17 context, for nearly all 32,768 variant 5’ss of the form NNN/GYNNNN. Data were split 60:20:20 into training, validation, and test sets. Latent phenotype models with one of four types of G-P map (additive, neighbor, pairwise, or black box), as well as a GE measurement process with a heteroscedastic skewed-t noise model, were inferred. The epistasis package of Sailer and Harms^24^ was also used to infer an additive G-P map and GE nonlinearity. (**a**) Performance of trained models as quantified by *I*_var_ and *I*_pre_, computed on test data. The lower bound on *I*_int_ was estimated from experimental replicates (see **Methods**). p-value reflects a two-sided z-test. *I*_var_ was not computed for the additive (epistasis package) model because that package does not infer an explicit noise model. (**b-d**) Measurement values versus latent phenotype values, computed on test data, using the additive (epistasis package) model (**b**), the additive model (**c**), and the pairwise model (**d**). The corresponding GE measurement processes are also shown. (**e**) Sequence logo^46^ illustrating the additive effects component of the pairwise G-P map. Dashed line indicates the exon/intron boundary. G at +1 serves as a placeholder because no other bases were assayed at this position. Only values for U and C at +2 were inferred. (**f**) Heatmap showing the pairwise effects component of the pairwise G-P map. White diagonals correspond to unobserved bases. Error bars indicate standard errors. **Supplemental Information Fig. S3** shows the uncertainties in these parameters. MPSA: massively parallel splicing assay; PSI: percent spliced in; G-P: genotype-phenotype; GE: global epistasis.

We also observe that increasingly complex G-P maps exhibit increased accuracy. For example, the additive G-P map gives *I*_pre_ = 0.257 ± 0.013 bits, whereas the pairwise G-P map (**Figs. 5e**,**f**) attains *I*_pre_ = 0.374 ± 0.014 bits. Using MAVE-NN’s built-in parametric bootstrap approach for quantifying parameter uncertainty, we find that both the additive and pairwise G-P map parameters are very precisely determined (see **Supplemental Information Fig. S3**). The black box G-P map, which is comprised of 5 densely connected hidden layers of 10 nodes each, performed the best of all four G-P maps, achieving *I*_pre_ = 0.458 ± 0.015 bits. Remarkably, this last predictive information value exceeds the lower bound of *I*_int_ ≥ 0.462 ± 0.009 bits, which was estimated from replicate experiments (see **Methods**). We thus conclude that pairwise interaction models are not flexible enough to fully account for how 5’ss sequences control splicing. More generally, these results underscore the need for software that is capable of inferring and assessing a variety of different G-P maps through a uniform interface.

### Application: biophysically interpretable G-P maps

Biophysical models, unlike the phenomenological models considered thus far, have mathematical structures that reflect specific hypotheses about how sequence-dependent interactions between macromolecules mechanistically define G-P maps. Thermodynamic models, which rely on a quasi-equilibrium assumption, are the most commonly used type of biophysical model.^39–41^ Previous studies have shown that precise thermodynamic models can be inferred from MAVE datasets,^16^ but no software intended for use by the broader MAVE community has yet been developed for doing this. MAVE-NN meets this need by enabling the inference of custom G-P maps. We now demonstrate this biophysical modeling capability in the contexts of protein-ligand binding (using DMS data; **Fig. 1a**) and bacterial transcriptional regulation (using sort-seq MPRA data; **Fig. 1e**). An expanded discussion of how these models are mathematically formulated and specified within MAVE-NN is provided in the “Biophysical modeling” section of **Supplemental Information**.

Otwinowski^42^ showed that a three-state thermodynamic G-P map (**Fig. 6a**), one that accounts for GB1 folding energy in addition to GB1-IgG binding energy,^43^ can explain the DMS data of Olson et al.^35^ better than a simple additive G-P map does. This biophysical model subsequently received impressive confirmation in the work of Nisthal et al.,^44^ who measured the thermostability of 812 single-mutation GB1 variants. We tested the ability of MAVE-NN to recover the same type of thermodynamic model that Otwinowski had inferred using custom analysis scripts. Our analysis yielded a G-P map with significantly improved performance on the data of Olson et al. (*I*_var_ = 2.303 ± 0.013 bits, *I*_pre_ = 2.357 ± 0.007 bits, *R*^2^ = 0.947 ± 0.001) relative to the additive G-P map of **Fig. 3. Fig. 6b** shows the two inferred energy matrices that respectively describe the effects of every possible single-residue mutation on the Gibbs free energies of protein folding and protein-ligand binding. The folding energy predictions of our model also correlate as well with the data of Nisthal et al. (*R*^2^ = 0.570 ± 0.049) as the predictions of Otwinowski’s model does (*R*^2^ = 0.515 ± 0.056). This demonstrates that MAVE-NN can infer accurate and interpretable quantitative models of protein biophysics.

**Figure 6.**
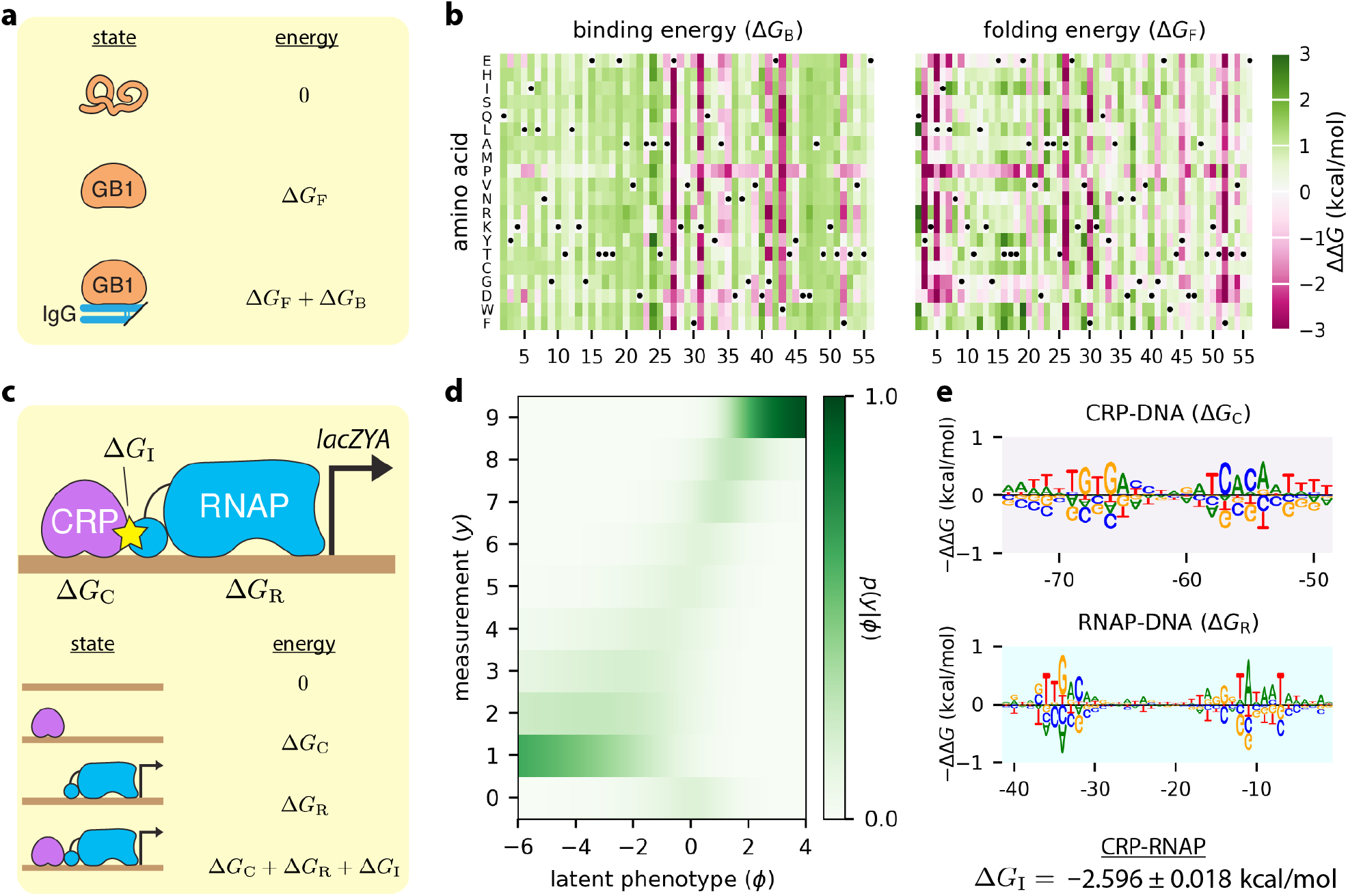
Biophysical models inferred from DMS and MPRA data. (**a**) Thermodynamic model for IgG binding by GB1. This model comprises three GB1 microstates (unfolded, folded-unbound, and folded-bound). The Gibbs free energies of folding (Δ*G*_*F*_) and binding (Δ*G*_*B*_) are computed from sequence using additive models called energy matrices. The latent phenotype is given by the fraction of time GB1 is in the folded-bound state. (**b**) The ΔΔ*G* parameters of the energy matrices for folding and binding, inferred from the data of Olson et al.^35^ using GE regression. **Supplemental Fig. S5** plots folding energy predictions against the measurements of Nisthal et al..^44^ (**c**) A four-state thermodynamic model for transcriptional activation at the *E. coli lac* promoter. The Gibbs free energies of RNAP-DNA binding (Δ*G*_R_) and CRP-DNA binding (Δ*G*_C_) are computed using energy matrices, whereas the CRP-RNAP interaction energy Δ*G*_I_ is a scalar. The latent phenotype is the fraction of time a promoter is bound by RNAP. (**d**,**e**) The latent phenotype model inferred from the sort-seq MPRA of Kinney et al.,^16^ including both the MPA measurement process (**d**) and the parameters of the thermodynamic G-P map (**e**). Supplemental **Fig. S4** provides detailed definitions of the thermodynamic models in panels **a** and **c**. Sequence logos in panel **e** were generated using Logomaker,^46^ and standard errors for the protein-protein interaction energy were determined using the built-in parametric bootstrapping approach described in Methods. GE: global epistasis. RNAP: RNA polymerase. MPA: measurement-process agnostic. G-P: genotype-phenotype.

To test MAVE-NN’s ability to infer thermodynamic models of transcriptional regulation, we re-analyzed the MPRA data of Kinney et al.,^16^ in which random mutations to a 75 bp region of the *Escherichia coli lac* promoter were assayed. This promoter region binds two regulatory proteins, σ^70^ RNA polymerase (RNAP) and the transcription factor CRP. As in Kinney et al.,^16^ we proposed a four-state thermodynamic model that quantitatively explains how promoter sequences control transcription rate (**Fig. 6c**). The parameters of this G-P map include the Gibbs free energy of interaction between CRP and RNAP (Δ*G*_I_), as well as energy matrices that describe the CRP-DNA and RNAP-DNA interaction energies. Because the sort-seq MPRA of Kinney et al. yielded discrete measurement values (**Figs. 1e**,**f**), we used an MPA measurement process in our latent phenotype model (**Fig. 6d**). The biophysical parameter values we thus inferred (**Fig. 6e**), including a CRP-RNAP interaction energy of Δ*G*_I_ = −2.598 ± 0.018 kcal/mol, largely match those of Kinney et al., but were obtained far more rapidly (in ∼10 min versus multiple days) thanks to the use of stochastic gradient descent rather than Metropolis Monte Carlo.

### Constraints on datasets and models

As stated above, MAVE-NN places certain limitations on both input datasets and latent phenotype models. Some of these constraints have been adopted to simplify the initial release of MAVE-NN and can potentially be relaxed in future updates. Others reflect fundamental mathematical properties of latent phenotype models. Here we summarize the primary constraints users should be aware of.

MAVE-NN currently requires that all input sequences be the same length. This constraint has been adopted because a large fraction of MAVE datasets have this form, and all of the built-in G-P maps operate only on fixed-length sequences. Users who wish to analyze variable length sequences can still do so by padding the ends of sequences with dummy characters. Alternatively, users can provide a multiple-sequence alignment as input and include the gap character as one of the characters to consider when training models.

As stated above, MAVE-NN can analyze MAVE datasets that have either continuous or discrete measurements. At present, both types of measurements must be one-dimensional, i.e., users cannot fit a single model to vectors of multiple measurements (e.g., joint measurements of protein binding affinity and protein stability, as in Ref. ^27^). This constraint has been adopted only to simplify the user interface of the initial release. It is not a fundamental limitation of latent phenotype models and is scheduled to be relaxed in upcoming versions of MAVE-NN.

The current implementation of MAVE-NN also supports only one-dimensional latent phenotypes (though the latent phenotype of custom G-P maps can depend on multiple precursor phenotypes, e.g., binding energy or folding energy). This restriction was made because accurately interpreting multi-dimensional latent phenotypes is substantially more fraught than interpreting one-dimensional latent phenotypes, and we believe that additional computational tools need to be developed to facilitate such interpretation. That being said, the mathematical form of latent phenotype models is fully compatible with multi-dimensional latent phenotypes. Indeed, this modeling strategy has been used in other work,^20,27,28,45^ and we plan to enable this functionality in future updates to MAVE-NN.

More fundamental constraints come into play when analyzing MAVE data that contains only single-mutation variants. In such experiments, the underlying effects of individual mutations are hopelessly confounded by the biophysical, physiological, and experimental nonlinearities that may be present. By contrast, when the same mutation is observed in multiple genetic backgrounds, MAVE-NN can use systematic differences in the mutational effects observed between stronger and weaker backgrounds to remove these confounding influences. Thus, for datasets that comprise only single-mutant effects, we limit MAVE-NN to inferring only additive G-P maps using GE regression, and while the noise model in the GE measurement process is allowed to be heteroscedastic, the nonlinearity is constrained to be linear.

We emphasize that, in practice, only a modest number of multiple-mutant variants are required for MAVE-NN to learn the form of a non-linear measurement process (see Fig. 3e-g). In this way, including a small fraction of the possible double-mutation variants in MAVE libraries can be beneficial even just for determining the effects of single mutations. Adding such non-comprehensive sets of double mutants to MAVE libraries is experimentally straight-forward, and our numerical experiments suggest that assaying roughly the same number of double-mutation variants as single-mutation variants should often suffice. We therefore recommend that experimentalists—even those primarily interested in the effects of single mutations—consider augmenting their MAVE libraries with a small subset of double-mutation variants.

## Discussion

In this work we have presented a unified strategy for inferring quantitative models of G-P maps from diverse MAVE datasets. At the core of our approach is the conceptualization of G-P maps as a form of information compression, i.e., the G-P map first compresses an input sequence into a latent phenotype value, which the MAVE then reads out indirectly via a noisy nonlinear measurement process. By explicitly modeling this measurement process, one can remove potentially confounding effects from the G-P map, as well as accommodate diverse experimental designs. We have also introduced three information-theoretic metrics for assessing the performance of the resulting models. These capabilities have been implemented within an easy-to-use Python package called MAVE-NN.

We have demonstrated the capabilities of MAVE-NN in diverse biological contexts, including in the analysis of both DMS and MPRA data. We have also demonstrated the superior performance of MAVE-NN relative to the epistasis package of Sailer and Harms.^24^ Along the way, we observed that MAVE-NN can deconvolve experimental nonlinearities from additive G-P maps when a relatively small number of sequences containing multiple mutations are included in the assayed libraries. This capability provides a compelling reason for experimentalists to include such sequences in their MAVE libraries, even if they are primarily interested in the effects of single mutations. Finally, we showed how MAVE-NN can learn biophysically interpretable G-P maps from both DMS and MPRA data.

MAVE-NN thus fills a critical need in the MAVE community, providing user-friendly software capable of learning quantitative models of G-P maps from diverse MAVE datasets. MAVE-NN has a streamlined user interface and is readily installed from PyPI by executing “pip install mavenn” at the command line. Comprehensive documentation and step-by-step tutorials are available at http://mavenn.readthedocs.io.

## Acknowledgements

This work was supported by NIH grant 1R35GM133777 (awarded to JBK), NIH Grant 1R35GM133613 (awarded to DMM), an Alfred P. Sloan Research Fellowship (awarded to DMM), a grant from the CSHL/Northwell Health partnership, and funding from the Simons Center for Quantitative Biology at Cold Spring Harbor Laboratory.

## Author Contributions

AT, WTI, DMM, and JBK conceived the project. AT and JBK wrote the software with assistance from AP and MK. WTI and JBK wrote a preliminary version of the software. AT, MK, and JBK performed the data analysis. AT, DMM, and JBK wrote the manuscript with contributions from MK and AP.

## Availability of Data and Materials

MAVE-NN can be installed from PyPI by executing “pip install mavenn” at the POSIX command line. Comprehensive documentation, including step-by-step tutorials, is available at http://mavenn.readthedocs.io. Source code, the data sets analyzed in this paper, and the scripts used for training the models and making the figures presented herein, are available under an MIT open-source license at https://github.com/jbkinney/mavenn. MAVE-NN version 1.0.0 was used for all of the analysis described in this manuscript and is archived on Zenodo at https://doi.org/10.5281/zenodo.5812439.

## Conflicts of Interest

The authors declare that they have no known conflicts of interest.

## Online Methods

### Notation

We represent each MAVE dataset as a set of *N* observations, 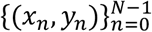, where each observation consists of a sequence *x*_*n*_ and a measurement *y*_*n*_. Here, *y*_*n*_ can be either a continuous real-valued number, or a nonnegative integer representing the “bin” in which the *n*th sequence was found. Note that, in this representation the same sequence *x* can be observed multiple times, potentially with different values for *y* due to experimental noise.

### G-P maps

We assume that all sequences have the same length *L*, and that at each of the *L* positions in each sequence there is one of *C* possible characters. MAVE-NN represents sequences using a vector of one-hot encoded features of the form

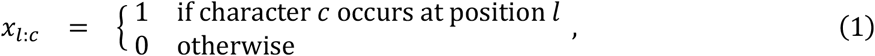

where *l* = 0,1, …, *L* − 1 indexes positions within the sequence, and *c* indexes the *C* distinct characters. MAVE-NN supports built-in alphabets for DNA, RNA and protein (with or without stop codons), as well as user-defined sequence alphabets.

We assume that the latent phenotype is given by a linear function *ϕ*(*x; θ*) that depends on a set of G-P map parameters *θ*. As mentioned in the main text, MAVE-NN supports four types of G-P map models, all of which can be inferred using either GE regression or MPA regression. The additive model is given by,

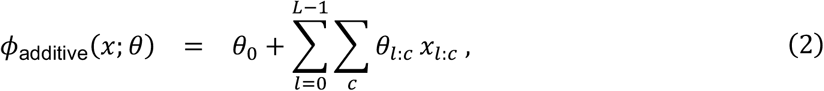

and thus each position in *x* contributes independently to the latent phenotype. The neighbor model is given by,

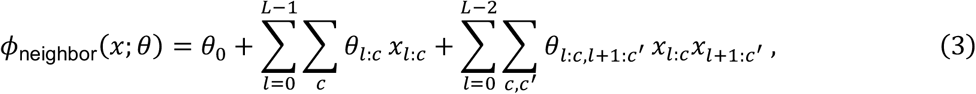

and further accounts for potential epistatic interactions between neighboring positions. The pairwise model is given by,

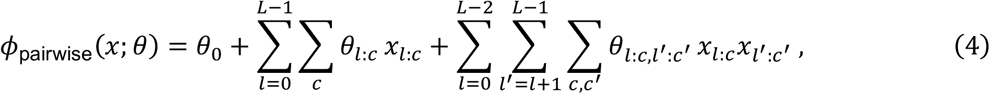

and includes interactions between all pairs of positions. Note our convention of requiring *l*′ > *l* in the pairwise parameters *θ*_*l:c,l*′ *:c*′_.

Unlike these three parametric models, the black box G-P map does not have a fixed functional form. Rather, it is given by a multilayer perceptron that takes a vector of sequence features (additive, neighbor, or pairwise) as input, contains multiple fully-connected hidden layers with nonlinear activations, and has a single node output with a linear activation. Users are able to specify the number of hidden layers, the number of nodes in each hidden layer, and the activation function used by these nodes.

MAVE-NN further supports custom G-P maps that users can define by subclassing the G-P map base class. These G-P maps can have arbitrary functional form, e.g., representing specific biophysical hypotheses of sequence function. This feature of MAVE-NN is showcased in the analyses of **Fig. 6**.

### Gauge modes and diffeomorphic modes

G-P maps typically have non-identifiable degrees of freedom that must be fixed, i.e., pinned down, before the values of individual parameters can be meaningfully interpreted or compared between models. These degrees of freedom come in two flavors: gauge modes and diffeomorphic modes. Gauge modes are changes to *θ* that do not alter the values of the latent phenotype *ϕ*. Diffeomorphic modes^15,20^ are changes to *θ* that do alter *ϕ*, but do so in ways that can be undone by transformations of the measurement process *p*(*y*|*ϕ*). As shown by Kinney and Atwal,^15,20^ the diffeomorphic modes of linear G-P maps like those considered here will in general correspond to affine transformations of *ϕ*, although additional unconstrained modes can occur in special situations.

MAVE-NN fixes both gauge modes and diffeomorphic modes of inferred models (except when using custom G-P maps). The diffeomorphic modes of G-P maps are fixed by transforming *θ* via

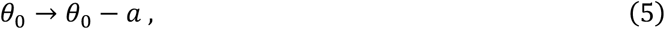

and then

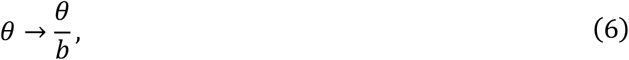

where *a* = mean({*ϕ*_*n*_}) and *b* = std({*ϕ*_*n*_}) are the mean and standard deviation of *ϕ* values computed on the training data. This produces a corresponding change in latent phenotype values *ϕ* → (*ϕ* − *a*)/*b*. To avoid altering likelihood values, MAVE-NN makes a corresponding transformation to the measurement process *p*(*y*|*ϕ*). In GE regression this is done by adjusting the GE nonlinearity via

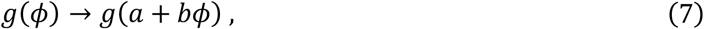

while keeping the noise model *p*(*y*|*ŷ*) fixed. In MPA regression MAVE-NN transforms the full measurement process via

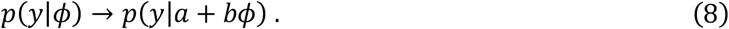

For the three parametric G-P maps, gauge modes are fixed using what we call the “hierarchical gauge.” Here, the parameters *θ* are adjusted so that the lower-order terms in *ϕ*(*x; θ*) account for the highest possible fraction of variance in *ϕ*. This procedure requires a probability distribution on sequence space with respect to which these variances are computed. MAVE-NN assumes that such distributions factorize by position, and can thus be represented by a probability matrix with elements *p*_*l:c*_, denoting the probability of character *c* at position *l*. MAVE-NN provides three built-in choices for this distribution: uniform, empirical, or wildtype. The corresponding values of *p*_*l:c*_ are given by

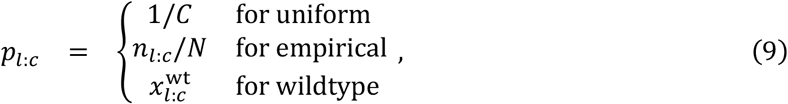

where *n*_*l:c*_ denotes the number of sequences (out of *N* total) that have character *c* at position *l*, and 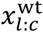 is the one-hot encoding of a user-specified wildtype sequence. In particular, the wildtype gauge was used for illustrating the additive G-P maps in **Fig. 3** and **Fig. 4**, while the uniform gauge was used for illustrating the pairwise G-P map in **Fig. 5** and the energy matrices in **Fig. 6**. After a sequence distribution is chosen, MAVE-NN fixes the gauge of the pairwise G-P map by transforming

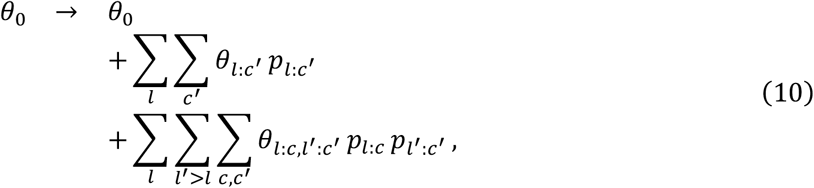

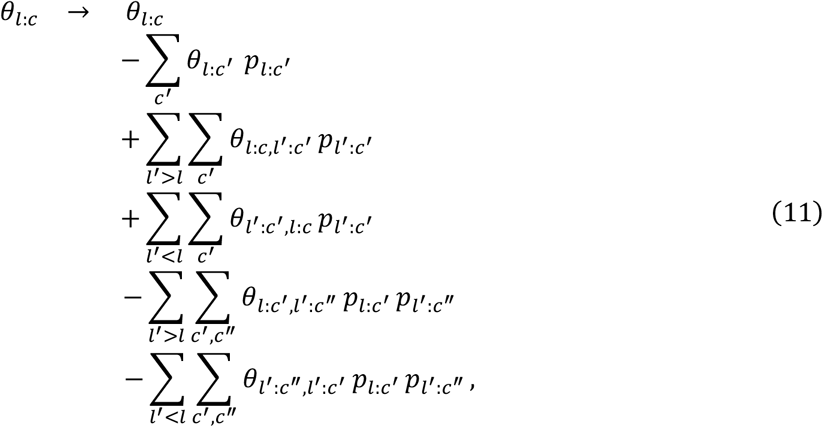

and

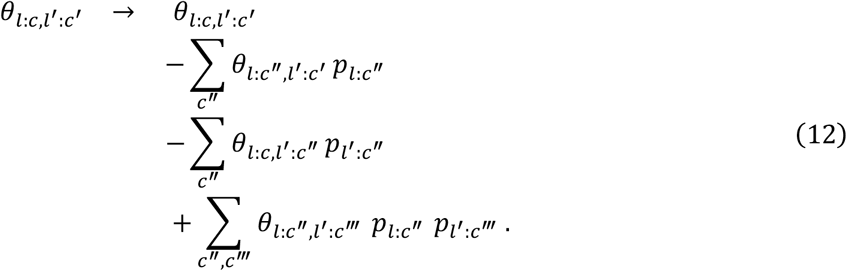

This transformation is also used for the additive and neighbor G-P maps, but with *θ*_*l:c,l*′ *:c*′_ = 0 for all *l, l*′ (additive) or whenever *l*′ ≠ *l* + 1 (neighbor).

### GE nonlinearities

GE models assume that each measurement *y* is a nonlinear function of the latent phenotype *g*(*ϕ*) plus some noise. In MAVE-NN, this nonlinearity is represented as a sum of tanh sigmoids:

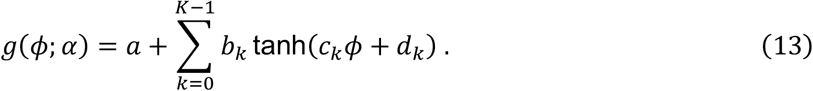

Here, *K* specifies the number of hidden nodes contributing to the sum, and *α* = {*a, b*_*k*_, *c*_*k*_, *d*_*k*_} are trainable parameters. We note that this mathematical form is an example of the bottleneck architecture previously used by^23,45^ for modeling GE nonlinearities. By default, MAVE-NN constrains *g*(*ϕ; α*) to be monotonic in *ϕ* by requiring all *b*_*k*_ ≥ 0 and *c*_*k*_ ≥ 0, but this constraint can be relaxed.

### GE noise models

MAVE-NN supports three types of GE noise model: Gaussian, Cauchy, and skew-t.

These all support the analytic computation of quantiles and confidence intervals, as well as the rapid sampling of simulated measurement values. The Gaussian noise model is given by

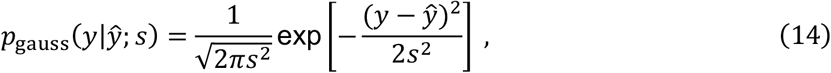

where *s* denotes the standard deviation. Importantly, MAVE-NN allows this noise model to be heteroskedastic by representing *s* as an exponentiated polynomial in *ŷ*, i.e.,

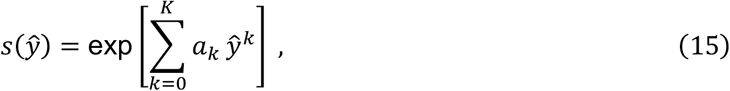

where *K* is the order of the polynomial and {*a*_*k*_} are trainable parameters. The user has the option to set *K*, and setting *K* = 0 renders this noise model homoscedastic. Quantiles are computed using *y*_*q*_ = *ŷ* + *s* √2 erf ^−1^(2*q* − 1) for user-specified values of *q* ∈ [0,1]. Similarly, the Cauchy noise model is given by

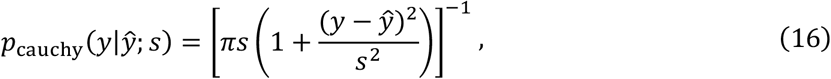

where the scale parameter *s* is an exponentiated *K*’th order polynomial in *ŷ*, and quantiles are computed using 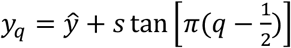.

The skew-t noise model is of the form described by Jones and Faddy,^29^ and is given by

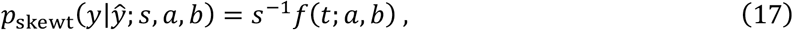

where

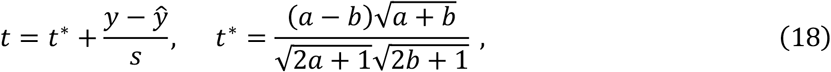

and

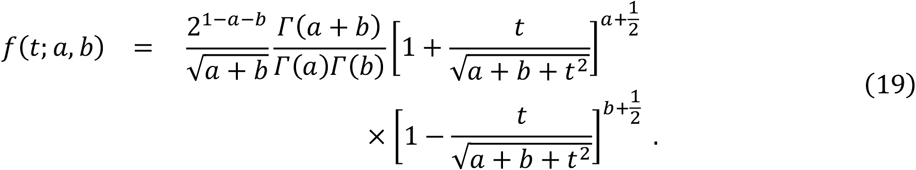

Note that the *t* statistic here is an affine function of *y* chosen so that the distribution’s mode (corresponding to *t*\*) is positioned at *ŷ*. The three parameters of this noise model, {*s, a, b*}, are each represented using *K*-th order exponentiated polynomials with trainable coefficients.

Quantiles are computed using

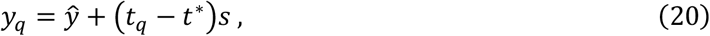

where

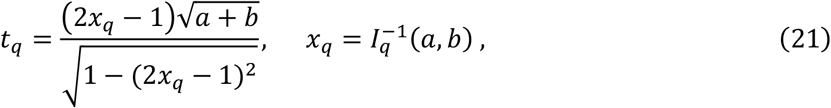

and *I*^−1^ denotes the inverse of the regularized incomplete Beta function *I*_*x*_(*a, b*).

### Empirical noise models

MAVE-NN further supports the inference of GE regression models that account for user-specified measurement noise. In such cases, the user provides a set of measurement-specific uncertainties 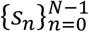 along with the corresponding sequences and measurements. These uncertainties can, for example, be estimated by using a software package like Enrich2^11^ or DiMSum^14^. MAVE-NN then trains the parameters of latent phenotype models by assuming a Gaussian noise model of the form

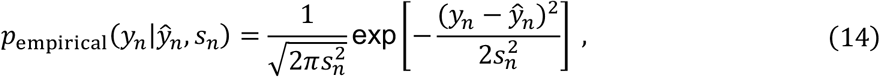

where *ŷ*_*n*_ = *g*(*f*(*x*_*n*_; *θ*); *α*) is the expected measurement for sequence *x*_*n*_, *θ* denotes G-P map parameters, and *α* denotes the parameters of the GE nonlinearity. This noise model thus has the advantage of having no free parameters, but it may be problematically mis-specified if the true error distribution is heavy-tailed or skewed.

### MPA measurement process

In MPA regression, MAVE-NN directly models the measurement process *p*(*y*|*ϕ*). At present, MAVE-NN only supports MPA regression for discrete values of *y* indexed using nonnegative integers. MAVE-NN supports two alternative forms of input for MPA regression. One is a set of sequence-measurement pairs 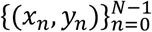, where *N* is the total number of reads, {*x*_*n*_} is a set of (typically) non-unique sequences, each *y*_*n*_ ∈ {0,1, …, *Y* − 1} is a bin number, and *Y* is the total number of bins. The other is a set of sequence-count-vector pairs 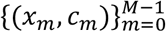, where *M* is the total number of unique sequences and *c*_*m*_ = (*c*_*m*0_, *c*_*m*1_, …, *c*_*m(Y*−1)_) is a vector that lists the number of times *c*_*my*_ that the sequence *x*_*m*_ was observed in each bin *y*. MPA measurement processes are represented as multilayer perceptron with one hidden layer (having tanh activations) and a softmax output layer. Specifically,

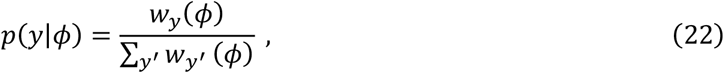

where

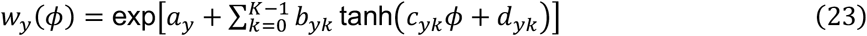

and *K* is the number of hidden nodes per value of *y*. The trainable parameters of this measurement process are *η* = {*a*_*y*_, *b*_*yk*_, *c*_*yk*_, *d*_*yk*_}.

### Loss function

Let *θ* denote the G-P map parameters, and *η* denote the parameters of the measurement process. MAVE-NN optimizes these parameters using stochastic gradient descent on a loss function given by

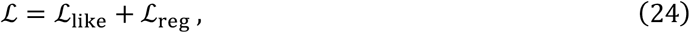

where ℒ_like_ is the negative log likelihood of the model, given by

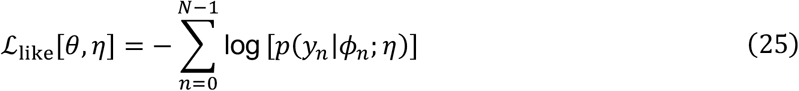

where *ϕ*_*n*_ = *ϕ*(*x*_*n*_; *θ*), and ℒ_reg_ provides for regularization of the model parameters.

In the context of GE regression, we can write *η* = (*α, β*) where *α* represents the parameters of the GE nonlinearity *g*(*ϕ; α*), and *β* denotes the parameters of the noise model *p*(*y*|*ŷ; β*). The likelihood contribution from each observation *n* then becomes *p*(*y*_*n*_|*ϕ*_*n*_; *η*) = *p*(*y*_*n*_|*ŷ*_*n*_; *β*) where *ŷ*_*n*_ = *g*(*ϕ*_*n*_; *α*). In the context of MPA regression with a dataset of the form 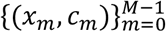, the loss function simplifies to

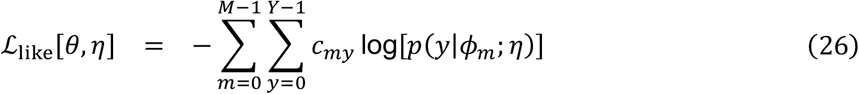

where *ϕ*_*m*_ = *ϕ*(*x*_*m*_; *θ*). For the regularization term, MAVE-NN uses an *L*_2_ penalty of the form

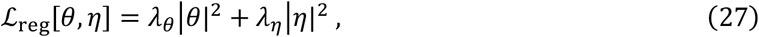

where the user-adjusted parameters *λ*_*θ*_ and *λ*_*η*_ respectively control the strength of regularization for the G-P map and measurement process parameters.

### Predictive information

In what follows, we use *p*_model_(*y*|*ϕ*) to denote a measurement process inferred by MAVE-NN, whereas *p*_true_(*y*|*ϕ*) denotes the empirical conditional distribution of *y* and *ϕ* values that would be observed in the limit of infinite test data.

Predictive information *I*_pre_ = *I*[*y; ϕ*], where *I*[·;·] represents mutual information computed on data not used for training (i.e., a held-out test set or data from a different experiment), *I*_pre_ provides a measure of how strongly a G-P map predicts experimental measurements. Importantly, this quantity does not depend on the corresponding measurement process *p*_model_(*y*|*ϕ*). To estimate *I*_pre_, we use k’th nearest neighbor (kNN) estimators of entropy and mutual information adapted from the NPEET Python package.^47^ Here, the user has the option of adjusting *k*, which controls a variance/bias tradeoff. When *y* is discrete (MPA regression), *I*_pre_ is computed using the classic kNN entropy estimator^48,49^ via the decomposition *I*[*y; ϕ*] = *H*[*ϕ*] − ∑_*y*_ *p* (*y*)*H*_*y*_[*ϕ*], where *H*_*y*_[*ϕ*] denotes the entropy of *p*_true_(*ϕ*|*y*). When *y* is continuous (GE regression), *I*[*y; ϕ*] is estimated using the kNN-based Kraskov Stögbauer Grassberger (KSG) algorithm.^49^ This approach optionally supports the local nonuniformity correction of Gao et al., which is important when *y* and *ϕ* exhibit strong dependencies, but which also requires substantially more time to compute.

### Variational information

We define variational information as an affine transformation of ℒ_like_,

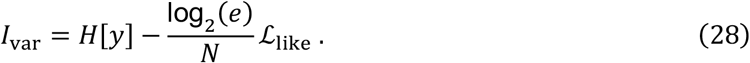

Here, *H*[*y*] is the entropy of the data {*y*_*n*_}, which is estimated using the *k*’th nearest neighbor (kNN) estimator from the NPEET package.^47^ Noting that this quantity can also be written as *I*_var_ = *H*[*y*] − mean({*Q*_*n*_}), where *Q*_*n*_ = −log_2_*p*(*y*_*n*_|*ϕ*_*n*_), we estimate the associated uncertainty using

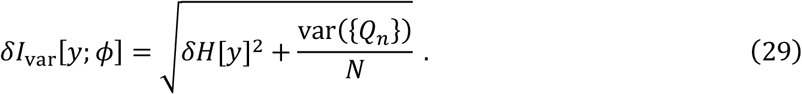

The inference strategy used by MAVE-NN is based on the fact that *I*_var_ provides a tight variational lower bound on *I*_pre_. Indeed, in the large data limit,

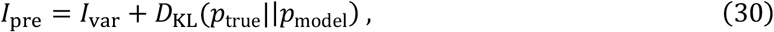

where *D*_KL_(·) ≥ 0 is the Kullback-Leibler divergence, and thus quantifies the accuracy of the inferred measurement process. From **Eq. 30** one can see that, with appropriate caveats, maximizing *I*_var_ (or equivalently, ℒ_like_) will also maximize *I*_pre_.^20^ But unlike *I*_pre_, *I*_var_ is readily compatible with backpropagation and stochastic gradient descent. See Supplemental Information for a derivation of **Eq. 30** and an expanded discussion of this key point. Note: Sharpee et al.^50^ cleverly showed that *I*_pre_ can, in fact, be optimized using stochastic gradient descent. Computing gradients of *I*_pre_, however, requires a time-consuming density estimation step. Optimizing *I*_var_, on the other hand, can be done using standard per-datum backpropagation.

### Intrinsic information

Intrinsic information, *I*_int_ = *I*[*x; y*], is the mutual information between the sequences *x* and measurements *y* in a dataset. This quantity is somewhat tricky to estimate due to the high-dimensional nature of sequence space. We instead used three different methods to obtain the upper and lower bounds on *I*_int_ shown in **Fig. 3d** and **Fig. 5a**. More generally, we believe the development of both computational and experimental methods for estimating *I*_int_ is be an important avenue for future research.

To compute the upper bound on *I*_int_ for GB1 data (in **Fig. 3d**), we used the fact that

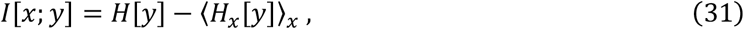

where *H*[*y*] is the entropy of all measurements *y, H*_*x*_[*y*] is the entropy of *p*(*y*|*x*) for a specific choice of sequence *x*, and ⟨·⟩_*x*_ indicates averaging over all sequences *x*. In this dataset, the measurement values were computed using

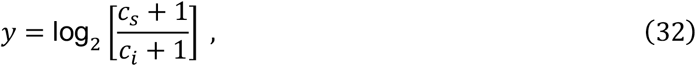

where *c*_*i*_ is the input read count and *c*_*s*_ is the selected read count. *H*[*y*] was estimated using the KNN estimator.^48^ We estimated the uncertainty in *y* by propagating errors expected due to Poisson fluctuations in read counts, which gives

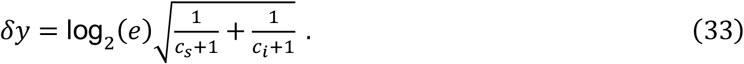

Then, assuming *p*(*y*|*x*) to be approximately Gaussian, we find the corresponding conditional entropy to be

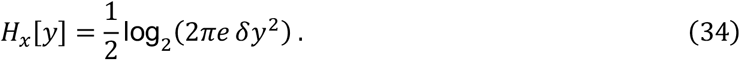

These *H*[*y*] and *H*_*x*_[*y*] values were then used in **Eq. 31** to estimate *I*_int_. This should provide an upper bound on the true value of *I*_int_ because uncertainty in *y* must be at least that expected under Poisson sampling of reads. We note, however, that the use of linear error propagation and the assumption that *p*(*y*|*x*) is approximately Gaussian complicate this conclusion. Also, when applied to MPSA data, this method yielded an upper bound of 0.96 bits. We believe this value is likely to be far higher than the true value of *I*_int_, and that this mismatch probably resulted from read counts in the MPSA data being over-dispersed.

To compute the lower bound on *I*_int_ for GB1 data (**Fig. 3d**) we used the predictive information *I*_pre_ (on test data) of a GE regression model having a blackbox G-P map. This provides a lower bound because *I*_int_ ≥ *I*_pre_ for any model (when evaluated on test data) due to the Data Processing Inequality and the Markov Chain nature of the dependencies *y* ← *x* → *ϕ* in **Fig. 2e**.^20,31^

To compute a lower bound on *I*_int_ for MPSA data (**Fig. *5*c**), we leveraged the availability of replicate data in Wong et al..^38^ Let *y* and *y*′ represent the original and replicate measurements obtained for a sequence *x*. Because *y* ← *x* → *y*′ forms a Markov chain, *I*[*x; y*] ≥ *I*[*y; y*′].^31^ We therefore used an estimate of *I*[*y; y*′], computed using the KSG method,^47,49^ as the lower bound for *I*_int_.

### Uncertainties in kNN estimates of mutual information

MAVE-NN quantifies uncertainties in *H*[*y*] and *I*[*y; ϕ*] using multiple random samples of half the data. Let 𝒟_100%_ denote a full dataset, and let 𝒟_50%,*r*_ denote a 50% subsample (indexed by *r*) of this dataset. Given an estimator *E*(·) of either entropy or mutual information, as well as the number of subsamples *R* to use, the uncertainty in *E*(𝒟_100%_) is estimated as

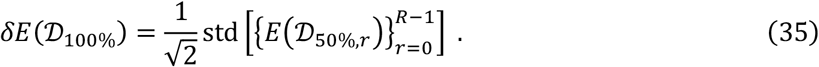

MAVE-NN uses *R* = 25 by default. We note that computing such uncertainty estimates substantially increases computation time, as *E*(·) needs to be evaluated *R* + 1 times instead of just once. We also note that bootstrap resampling^51,52^ is often inadvisable in this context, as it systematically underestimates *H*[*y*] and overestimates *I*[*y; z*].

### Uncertainties in G-P map parameters

Given a trained latent phenotype model, having G-P map parameters *θ*\* and measurement process parameters *η*\*, MAVE-NN can optionally assess model uncertainty using the following parametric bootstrap approach. Using the trained model with parameters (*θ*\*, *η*\*) as “ground truth”, MAVE-NN simulates *R* (chosen by the user) different MAVE datasets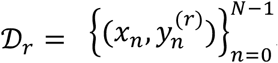, where *r* = 0,1, …, *R* − 1 indexes the different simulations. Note that the sequences in these simulated datasets are the same as the original training sequence, but the measurements are different. For each simulated dataset 𝒟_*r*_, MAVE-NN then trains a new model, by default using the same hyperparameters as were used for the ground truth model. This procedure yields a set 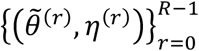 of simulation-inferred G-P map parameters and corresponding measurement process parameters. Users can then use this sampling of G-P map parameters to estimate uncertainties, e.g., by reporting 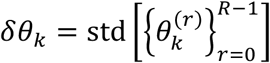.

An important detail when assessing parameter uncertainty is to ensure that both the gauge modes and the diffeomorphic modes of each model are fixed. This is necessary so that differences in the parameters that cannot result in changes to model predictions do not inflate the uncertainty estimates. For additive, neighbor, and pairwise G-P maps, MAVE-NN automatically implements the procedure described in the “Gauge modes and diffeomorphic modes” section above, thereby removing these extra degrees of freedom. However, for more complex models such as those implemented by MAVE-NN’s custom G-P map functionality (e.g., representing biophysical models) different gauge freedoms and diffeomorphic modes may arise depending on the details of the model, and users must take care to determine and fix these prior to assessing parameter uncertainty. We also note that no meaningful computation of individual parameter uncertainties is likely to be possible for highly over-parameterized models, such as the “black box” deep neural network models supported by MAVE-NN.

### Datasets

For the GB1 DMS dataset of Olson et al.,^35^ measurements were computed using

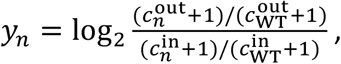

where 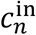 and 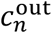 respectively represent the number of reads from the input and output samples (i.e., pre-selection and post-selection libraries), and *n* = WT represents the 55 aa wildtype sequence, corresponding to positions 2-56 of the GB1 domain. To infer the model in **Fig. 3b** and to compute the information metrics in **Fig. 3c**, only double-mutant sequences with 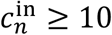 were used; these represent 530,737 out of the 536,085 possible double mutants. For the models in **Figs. 3d-f**, *y*_*n*_ values for the 1045 single-mutant were also used in the inference procedure.

For the Aβ DMS data of Seuma et al.^36^ and TDP-43 DMS data of Bolognesi et al.,^37^ *y*_*n*_ values respectively represent nucleation scores and toxicity scores reported by the authors.

For the MPSA data of Wong et al.,^38^ we used the data of library 1 replicate 1 obtained for the *BRCA2* minigene data. Measurements were computed as

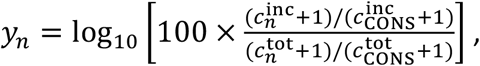

where 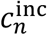 and 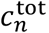 respectively represent the number of barcode reads obtained from exon inclusion isoforms and from total mRNA, and *n* = CONS corresponds to the consensus 5’ss sequence CAG/GUAAGU. Corresponding PSI values were computed as 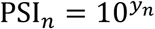. Only sequences with 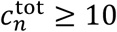 were used, representing 30,483 of the 32,768 possible sequences of the form NNN/GYNNNN.

For the *lac* promoter sort-seq MPRA data of Kinney et al.,^16^ we used data from the “full-wt” experiment (available at https://github.com/jbkinney/09_sortseq).

## Notes

https://mavenn.readthedocs.io/

### Competing Interest Statement

The authors have declared no competing interest.

### Summary of Updates

Moderate revisions throughout.

## References

1. Kinney, J. B. & McCandlish, D. M. Massively parallel assays and quantitative sequence-function relationships. Annu Rev Genom Hum G 20, 99–127 (2019).

2. Starita, L. M. et al. Variant interpretation: functional assays to the rescue. Am J Hum Genetics 101, 315–325 (2017).

3. Fowler, D. M. & Fields, S. Deep mutational scanning: a new style of protein science. Nat Methods 11, 801–807 (2014).

4. Levo, M. & Segal, E. In pursuit of design principles of regulatory sequences. Nat Rev Genet 15, 453–468 (2014).

5. White, M. A. Understanding how cis-regulatory function is encoded in DNA sequence using massively parallel reporter assays and designed sequences. Genomics 106, 165–170 (2015).

6. Inoue, F. & Ahituv, N. Decoding enhancers using massively parallel reporter assays. Genomics 106, 159–164 (2015).

7. Peterman, N. & Levine, E. Sort-seq under the hood: implications of design choices on large-scale characterization of sequence-function relations. BMC Genomics 17, 206 (2016).

8. Fowler, D. M., Araya, C. L., Gerard, W. & Fields, S. Enrich: software for analysis of protein function by enrichment and depletion of variants. Bioinformatics 27, 3430–3431 (2011).

9. Alam, K. K., Chang, J. L. & Burke, D. H. FASTAptamer: A bioinformatic toolkit for high-throughput sequence analysis of combinatorial selections. Mol Ther-Nucleic Acids 4, e230 (2015).

10. Bloom, J. D. Software for the analysis and visualization of deep mutational scanning data. BMC Bioinformatics 16, 168 (2015).

11. Rubin, A. F. et al. A statistical framework for analyzing deep mutational scanning data. Genome Biol 18, 1–15 (2017).

12. Ashuach, T. et al. MPRAnalyze: statistical framework for massively parallel reporter assays. Genome Biol 20, 183 (2019).

13. Niroula, A., Ajore, R. & Nilsson, B. MPRAscore: robust and non-parametric analysis of massively parallel reporter assays. Bioinformatics 35, 5351–5353 (2019).

14. Faure, A. J., Schmiedel, J. M., Baeza-Centurion, P. & Lehner, B. DiMSum: an error model and pipeline for analyzing deep mutational scanning data and diagnosing common experimental pathologies. Genome Biol 21, 207 (2020).

15. Atwal, G. S. & Kinney, J. B. Learning quantitative sequence-function relationships from massively parallel experiments. J Stat Phys 162, 1203–1243 (2016).

16. Kinney, J. B., Murugan, A., Callan, C. G. & Cox, E. C. Using deep sequencing to characterize the biophysical mechanism of a transcriptional regulatory sequence. Proc Natl Acad Sci USA 107, 9158–9163 (2010).

17. Melnikov, A. et al. Systematic dissection and optimization of inducible enhancers in human cells using a massively parallel reporter assay. Nat Biotechnol 30, 271–277 (2012).

18. Mogno, I., Kwasnieski, J. C. & Cohen, B. A. Massively parallel synthetic promoter assays reveal the in vivo effects of binding site variants. Genome Res 23, 1908–1915 (2013).

19. Abadi, M. et al. TensorFlow: A Systems for Large-Scale Machine Learning. in Proceedings of the 12th USENIX Symposium on Operating Systems Design and Implementation (OSDI ‘16) (2016).

20. Kinney, J. B. & Atwal, G. S. Parametric inference in the large data limit using maximally informative models. Neural Comput 26, 637–653 (2014).

21. Kinney, J. B., Tkačik, G. & Callan, C. G. Precise physical models of protein-DNA interaction from high-throughput data. Proc Natl Acad Sci USA 104, 501–506 (2007).

22. Otwinowski, J. & Nemenman, I. Genotype to phenotype mapping and the fitness landscape of the E. coli lac promoter. PLoS ONE 8, e61570 (2013).

23. Sarkisyan, K. S. et al. Local fitness landscape of the green fluorescent protein. Nature 533, 397–401 (2016).

24. Sailer, Z. R. & Harms, M. J. Detecting high-order epistasis in nonlinear genotype-phenotype maps. Genetics 205, 1079–1088 (2017).

25. Otwinowski, J., McCandlish, D. M. & Plotkin, J. B. Inferring the shape of global epistasis. Proc Natl Acad Sci USA 115, E7550–E7558 (2018).

26. Gelman, S., Fahlberg, S. A., Heinzelman, P., Romero, P. A. & Gitter, A. Neural networks to learn protein sequence-function relationships from deep mutational scanning data. Proc Natl Acad Sci USA 118, e2104878118 (2021).

27. Faure, A. J. et al. Global mapping of the energetic and allosteric landscapes of protein binding domains. bioRxiv doi:10.1101/2021.09.14.460249 (2021)

28. Tonner, P. D., Pressman, A. & Ross, D. Interpretable modeling of genotype-phenotype landscapes with state-of-the-art predictive power. bioRxiv doi:10.1101/2021.06.11.448129 (2021).

29. Jones, M. C. & Faddy, M. J. A skew extension of the t-distribution, with applications. J Roy Stat Soc B 65, 159–174 (2003).

30. Kinney, J. B. & Atwal, G. S. Equitability, mutual information, and the maximal information coefficient. Proc Natl Acad Sci USA 111, 3354–3359 (2014).

31. Cover, T. M. & Thomas, J. A. Elements of information theory. (Wiley, 2006).

32. Barber, D. & Agakov, F. The IM algorithm: a variational approach to information maximization. Advances in neural information processing systems 16. (2004).

33. Alemi, A. A., Fischer, I., Dillon, J. V. & Murphy, K. Deep variational information bottleneck. 1612.00410 (2016).

34. Chalk, M., Marre, O. & Tkačik, G. Relevant sparse codes with variational information bottleneck. arXiv:1605.07332 (2016).

35. Olson, C. A., Wu, N. C. & Sun, R. A comprehensive biophysical description of pairwise epistasis throughout an entire protein domain. Curr Biol 24, 2643–2651 (2014).

36. Seuma, M., Faure, A., Badia, M., Lehner, B. & Bolognesi, B. The genetic landscape for amyloid beta fibril nucleation accurately discriminates familial Alzheimer’s disease mutations. eLife 10, e63364 (2021).

37. Bolognesi, B. et al. The mutational landscape of a prion-like domain. Nat Commun 10, 4162 (2019).

38. Wong, M. S., Kinney, J. B. & Krainer, A. R. Quantitative activity profile and context dependence of all human 5’ splice sites. Mol Cell 71, 1012-1026.e3 (2018).

39. Bintu, L. et al. Transcriptional regulation by the numbers: models. Curr Opin Genet Dev 15, 116–124 (2005).

40. Sherman, M. S. & Cohen, B. A. Thermodynamic state ensemble models of cis-regulation. PLoS Comput Biol 8, e1002407 (2012).

41. Wong, F. & Gunawardena, J. Gene Regulation in and out of equilibrium. Annu Rev Biophys 49, 199–226 (2020).

42. Otwinowski, J. Biophysical inference of epistasis and the effects of mutations on protein stability and function. Mol Biol Evol 35, 2345–2354 (2018).

43. Manhart, M. & Morozov, A. V. Protein folding and binding can emerge as evolutionary spandrels through structural coupling. Proc Natl Acad Sci USA 112, 1797–1802 (2015).

44. Nisthal, A., Wang, C. Y., Ary, M. L. & Mayo, S. L. Protein stability engineering insights revealed by domain-wide comprehensive mutagenesis. Proc Natl Acad Sci USA 116, 16367–16377 (2019).

45. Pokusaeva, V. O. et al. An experimental assay of the interactions of amino acids from orthologous sequences shaping a complex fitness landscape. PLoS Genet 15, e1008079 (2019).

46. Tareen, A. & Kinney, J. B. Logomaker: beautiful sequence logos in Python. Bioinformatics 36, 2272–2274 (2020).

47. Steeg, G. V. Non-Parametric Entropy Estimation Toolbox (NPEET). https://www.isi.edu/~gregv/npeet.html (2014).

48. Vasicek, O. A test for normality based on sample entropy. J Roy Stat Soc B 38, 54–59 (1976).

49. Kraskov, A., Stögbauer, H. & Grassberger, P. Estimating mutual information. Phys Rev E 69, 066138 (2004).

50. Sharpee, T., Rust, N. C. & Bialek, W. Analyzing neural responses to natural signals: maximally informative dimensions. Neural Comput 16, 223–250 (2004).

51. Efron, B. Bootstrap methods: another look at the jackknife. Ann Stat 7, 1–26 (1979)..

52. Efron, B. & Tibshirani, R. Bootstrap methods for standard errors, confidence intervals, and other measures of statistical accuracy. Stat Sci 1, 54–75 (1986).

