## Supplemental Information for "MAVE-NN: learning genotype-phenotype maps from multiplex assays of variant effect"

**Justin B. Kinney**

### S1 Variational information as a lower bound on predictive information

Here we show that $I_{\text{var}}$ provides a variational lower bound to $I_{\text{pre}} = I[y;\phi]$. This fact was previously described by [S1] in the general context of mutual information maximization, then later discussed in the context of MAVE data analysis by [S2, S3]. Notably, it also plays an important role in the variational information bottleneck [S4, S5]. To our knowledge, however, $I_{\text{var}}$ itself has not previously been advocated as a useful information-like metric.

As in the main text, let $p_{\text{true}}(y,\phi)$ denote the joint density of $y$ and model-assigned $\phi$ values that would be observed on test data in the $N_{\text{test}} \to \infty$ limit, let $p_{\text{true}}(y|\phi)$ denote the corresponding conditional density, and let $p_{\text{model}}(y|\phi)$ denote the inferred measurement process of a model in question. We then recover Eq. 30 in Methods as follows:

$$
\begin{aligned}
I_{\text{pre}} &= H[y] - H[y|\phi] && \text{(S1)}\\
&= H[y] + \langle \log_2 p_{\text{true}}(y|\phi) \rangle_{\text{true}} && \text{(S2)}\\
&= H[y] + \langle \log_2 p_{\text{model}}(y|\phi) \rangle_{\text{true}} + \left\langle \log_2 \frac{p_{\text{true}}(y|\phi)}{p_{\text{model}}(y|\phi)} \right\rangle_{\text{true}} \\
&= H[y] - \frac{\log_2(e)}{N_{\text{test}}} \mathcal{L}_{\text{like}} + D_{\text{KL}}(p_{\text{true}}||p_{\text{model}}) \\
&= I_{\text{var}} + D_{\text{KL}}(p_{\text{true}}||p_{\text{model}}) && \text{(S3)}
\end{aligned}
$$

where $D_{\text{KL}}$ denotes Kullback-Leibler divergence, and $\langle\cdot\rangle_{\text{true}}$ indicates averaging over $p_{\text{true}}(y,\phi)$.

Now consider the infinite training data limit. Also assume that the correct G-P map and measurement process are within the class of models considered, as is often the case when analyzing simulated data. Then it is simple to see that maximizing $\mathcal{L}_{\text{like}}$ (or equivalently, $I_{\text{var}}$) will push $D_{\text{KL}} \to 0$ and $I_{\text{pre}} \to I_{\text{int}}$. For such models, we therefore recover $I_{\text{var}} = I_{\text{pre}} = I_{\text{int}}$.

On finite datasets we must use approximate methods to estimate these information values. The inequalities $I_{\text{var}} \leq I_{\text{pre}} \leq I_{\text{true}}$ thus become approximate. Still, the uncertainties in these information quantities are often quite small due to the large size of typical MAVE datasets, and these inequalities serve well to guide judgements about model accuracy and completeness.

### S2 Analysis of simulated data

Here we assess the performance of MAVE-NN on realistic simulated DMS and MPSA datasets, thus demonstrating its ability to accurately recover latent phenotype parameters and measurement nonlinearities.

#### S2.1 Additive GE regression on simulated DMS data

We simulated a DMS dataset consisting of 530,737 GB1 protein sequences by using MAVE-NN's `simulate_dataset` method. This method simulates sequences according to a probability matrix $p_{l:c}$ that lists the probability of each character $c$ at each position $l$. For this simulation, $p_{l:c}$ was obtained from a model trained by MAVE-NN using GB1 data from [S6], as described in the main text. Measurements for simulated sequences were generated using the G-P map and measurement process from the same model. We then used MAVE-NN to train a separate GE regression model with an additive G-P map from these simulated data.

Fig. S1a shows simulated measurements $y$ vs. inferred phenotype values $\phi$. The inferred function $g(\cdot)$ shows remarkable agreement with the true function $g$, which is also shown on the same panel (Fig. S1a). Model predictions on held-out test data yielded a very high $R^2$ value with true model predictions (Fig. S1b). Plotting inferred latent phenotype values against true latent phenotype values also show near-perfect correspondence (Fig. S1c). A scatter plot of inferred G-P map parameters vs. true G-P map parameters also shows very high correspondence (Fig. S1d).

We also performed GE regression on datasets of similar composition having variable sizes. Model inference time on a standard laptop computer (3.1 GHz CPU) was recorded and is plotted vs. dataset size in Fig. S1e. Model inference for the original published dataset was fast and took $\sim$ 3 minutes. Additionally, model performance improved with increasing simulated dataset size, but even moderately sized datasets, e.g. 10,000 simulated observations, achieved very high $R^2$ values (Fig. S1f).

#### S2.2 Pairwise GE regression on simulated MPSA data

We generated a simulated dataset consisting of 30,483 $5'$ss sequences by using MAVE-NN's `simulate_dataset` method. This method simulates sequences according to a probability matrix $p_{l:c}$ that lists the probability of each character $c$ at each position $l$. $p_{l:c}$ was computed using MPSA data from [S7] using MAVE-NN. These simulated sequences represent a library based on the template sequence NNN/GYNNNN, where position +1 is fixed to have G, position +2 is fixed to have Y (meaning C or U equally likely), and with all other positions having N (meaning A,C,G, or U equally likely). Measurements for simulated sequences were generated using the pairwise model trained on MPSA data, described in Figs. 5d-f of the main text. We then used MAVE-NN to train a separate GE regression model with pairwise G-P map from these simulated data.

Fig. S3a shows simulated measurements $y$ vs. latent phenotype values $\phi$. The inferred GE nonlinearity shows remarkable agreement with the true GE nonlinearity. Inferred model predictions for held-out test-set sequences yielded a very high $R^2$ value with the true model predictions (Fig. S3b). Plotting inferred latent phenotype values against true latent phenotype values also shows near-perfect correspondence (Fig. S3c). A scatter plot of inferred G-P map parameters $\theta$ vs. true G-P map parameters also shows very high correspondence (Fig. S3d).

We also performed pairwise GE regression on datasets of similar composition having variable sizes. Model inference time on a standard laptop computer (3.1 GHz CPU) was recorded and is plotted vs. dataset size in Fig. S3e. Model inference for the original published dataset was fast and took $\sim$ 1 minute. Additionally, model performance improved with increasing simulated dataset size, but even moderately sized datasets, e.g. 10,000 simulated observations, achieved very high $R^2$ values (Fig. S3f).

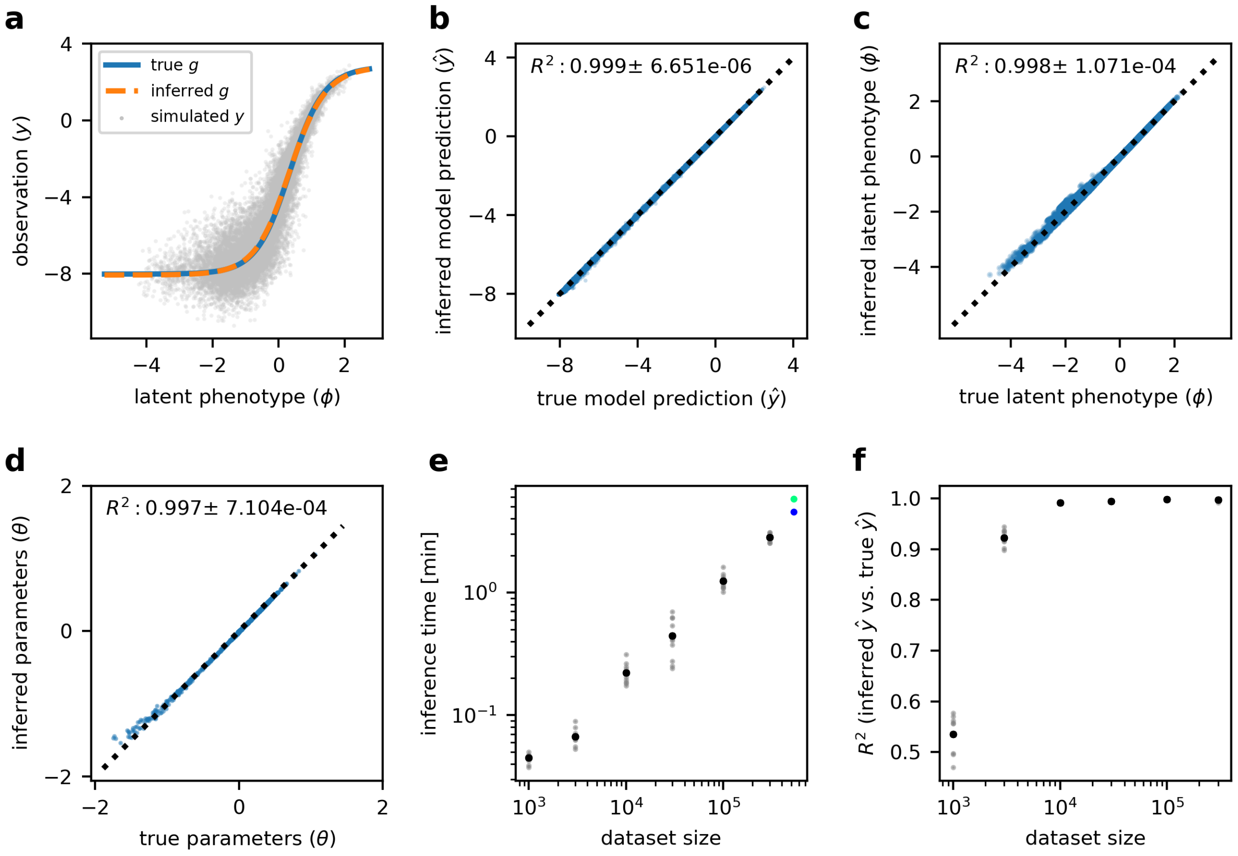


Figure S1: Additive GE regression on simulated GB1 data. A library of 530,737 simulated GB1 sequences was generated using MAVE-NN's `simulate_dataset` method, which used the GE model (true model) trained on the GB1 data from [S6] and illustrated in Figs. 3a,b of the main text. MAVE-NN was then used to train a separate GE regression model (inferred model) from these simulated data. (a) The true GE nonlinearity (blue curve) and simulated observation values $y$ (gray dots) are plotted, along with the inferred GE nonlinearity (dashed curve) against latent phenotype values $\phi$. (b) The inferred GE model achieves $R^2 = 0.997$ against true model predictions on held-out test data. (c) Plotting the inferred latent phenotype against the true latent phenotype shows near-perfect correspondence. (d) Inferred model parameters vs. true model parameters, again showing very high correspondence. (e) Inference time for datasets of varying size simulated in the same manner as above. The inference time for the real GB1 dataset is shown by the blue dot, while the inference time for the simulated library used to make panels (a-d) is indicated by the green dot. (f) $R^2$ values, as in panel b, as a function of dataset size. For each dataset size in (e-f), the black dot shows the mean inference time and mean $R^2$, respectively, for 10 simulations, whereas gray dots show individual values. For every simulation, we used the same network architecture and the same training, validation, and test dataset split as in the main text.

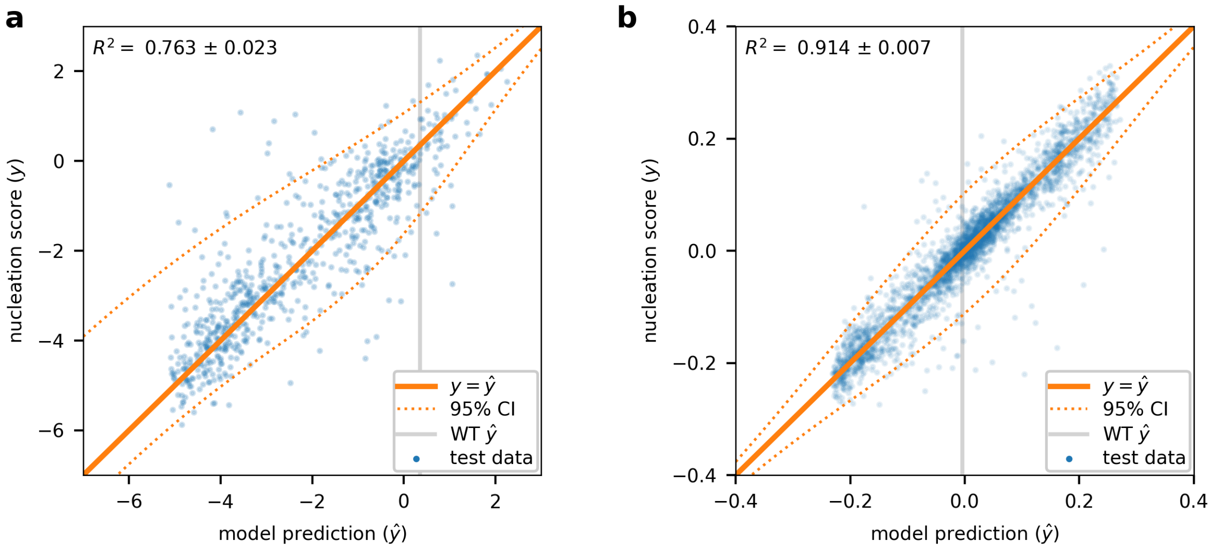


Figure S2: Additional analysis of DMS data on Aβ and TDP-43. **(a)** Nucleation scores vs. GE model predictions for Aβ. **(b)** Toxicity scores vs. GE model predictions predictions for TDP-43.

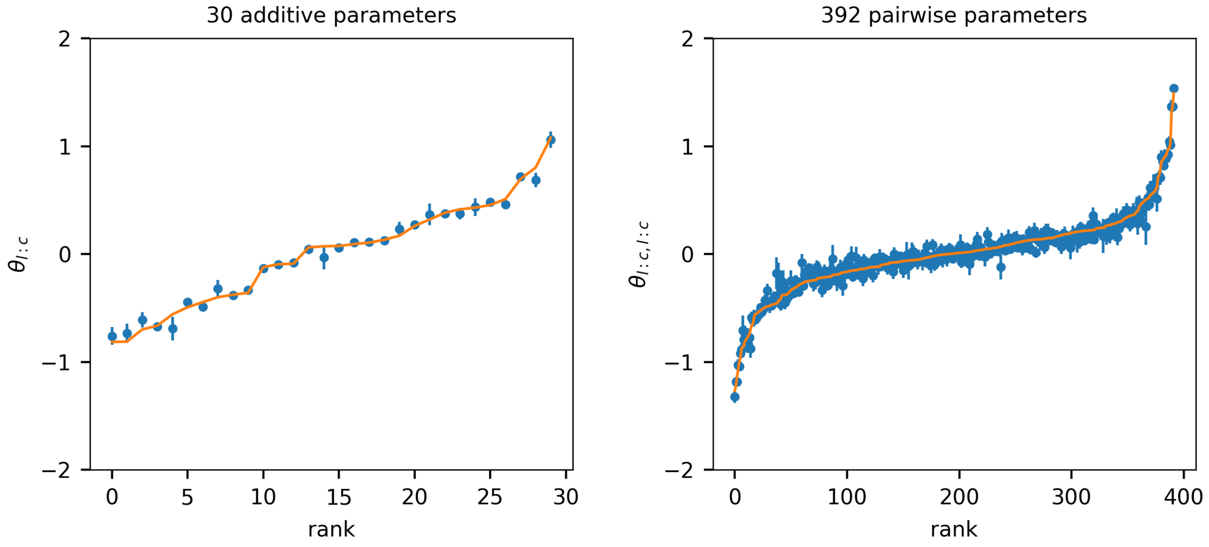


Figure S3: Parameter uncertainties inferred using MAVE-NN's parametric bootstrap functionality. Shown are the mean values (blue dots) and standard deviations (blue lines) of both the additive parameters (left panel) and pairwise parameters (right panel) of the pairwise G-P map inferred from the splicing MPRA data of Wong et al. [S7]. In each panel, parameters are arrayed along the horizontal axis according to the rank order of their best-fit values (orange line). Note that the values of these best-fit parameters are respectively illustrated in Figs. 5e and 5f of the main text.

### S3 Biophysical modeling using MAVE-NN

Fig. 6 of the main text illustrates three biophysical models trained as G-P maps using MAVE-NN. Here we review the general rationale for models of this form. We then describe the specific mathematical forms of the two thermodynamic models featured in Fig. 6. Please refer to the online documentation [S8] for details about how the equations presented below were coded within the MAVE-NN API to define custom G-P maps.

#### S3.1 General form of thermodynamic models

All of the biophysical models featured in Fig. 6 are examples of thermodynamic models. Thermodynamic models are defined by three key assumptions:

1. At each instant in time, the system of interest can occupy one of a fixed number of possible states.
2. The probability of the system being in each state depends on that state's Gibbs free energy via Boltzmann's law, which describes the probability one would observe for a system in thermal equilibrium.
3. The molecular phenotype of interest is a probability-weighted average of the activities of each individual state.

Mathematically, this means that the latent phenotype $\phi(x)$, for a given sequence $x$, is a weighted sum of state-specific activities:

$$\phi(x) = \sum_s \phi_s(x) p(s|x), \tag{S4}$$

where $s$ indexes possible states of the system, $\phi_s(x)$ is the (generally sequence-dependent) activity of state $s$, and $p(s|x)$ is the Boltzmann probability of state $s$. By Botlzmann's law, this probability is given by

$$p(s|x) = \frac{1}{Z(x)} \exp\left[-G_s(x)/k_B T\right], \tag{S5}$$

where $G_s(x)$ is the (generally sequence-dependent) Gibbs free energy of state $s$,

$$Z(x) = \sum_s \exp\left[-G_s(x)/k_B T\right] \tag{S6}$$

is a normalization factor that ensures the probabilities of all states sum to one, $k_B = 1.987 \times 10^{-3} \frac{\text{kcal}}{\text{mol}\cdot\text{K}}$ is Boltzmann's constant (which for our purposes is the same as the gas constant $R$), and $T$ is temperature in kelvin. Note that expressing model parameters in units of kcal/mol thus requires knowing the temperature at which the latent phenotype was measured: e.g., $k_B T = 0.582 \frac{\text{kcal}}{\text{mol}}$ at 20 °C (room temperature), while $k_B T = 0.616 \frac{\text{kcal}}{\text{mol}}$ at 37 °C (body temperature).

To infer biophysical models from MAVE data, one must propose specific mathematical formulas for how the state activities $\phi_s(x)$ and Gibbs free energies $G_s(x)$ depend on sequence. These quantities will depend on some set of *a priori* unknown parameters. These are the G-P map parameters $\theta$. The mathematical form of the G-P map itself, $\phi(x;\theta)$, then follows from Eqs. S4, S5, and S6.

To use a biophysical model as a G-P map with MAVE-NN, users must provide the specific equation for $\phi(x;\theta)$, written in terms of a one-hot encoded sequence vector $\vec{x}$, to the MAVE-NN API through subclassing. Please refer to the online documentation [S8] for details on how to do this.

#### S3.2 Thermodynamic model for protein GB1

Fig. S4a illustrates the thermodynamic model for protein GB1 that was proposed by [S9] to explain the DMS data of [S6], and which is featured in Figs. 6a and 6b of the main text. This model assumes three possible states for GB1: (1) unfolded, (2) folded but not bound to IgG, and (3) folded and bound to IgG. The activity in question, $\phi(x)$, is the fraction of time a GB1 molecule with peptide sequence $x$ is bound to IgG. The corresponding state-specific activities are $\phi_{(1)} = 0$, $\phi_{(2)} = 0$, and $\phi_{(3)} = 1$. Note that, both here and in the models below, we assume that all $\phi_s$ are fixed binary numbers that are known *a priori*, i.e., they do not depend on sequence $x$ or on any trainable parameters in $\theta$.

The unfolded state (1) is taken to be the reference state and thus assigned a Gibbs free energy of 0. $\Delta G_\text{F}$ denotes the Gibbs free energy of the folded state relative to the unfolded state, and thus state (2) has energy $\Delta G_\text{F}$. $\Delta G_\text{B}$ denotes the Gibbs free energy change upon the binding of a folded GB1 molecule to IgG, and thus the energy of state (3) is $\Delta G_\text{F} + \Delta G_\text{B}$. Each of these energies is assumed to be an additive function of peptide sequence $x$; the mathematical formulas for $\Delta G_\text{F}$ and $\Delta G_\text{B}$ in terms of the one-hot encoded vector $\vec{x}$ are shown. The parameters that control these two energies are $\vec{\theta}_\text{F}$, $\theta^0_\text{F}$ (for $\Delta G_\text{F}$) and $\vec{\theta}_\text{B}$, $\theta^0_\text{B}$ (for $\Delta G_\text{B}$). The resulting set of trainable G-P map parameters is

$$\theta = \left\{ \theta^0_\text{F}, \vec{\theta}_\text{F}, \theta^0_\text{B}, \vec{\theta}_\text{B} \right\}. \tag{S7}$$

To infer values for these parameters using MAVE-NN, we paired the biophysical G-P map $\phi(x;\theta)$ with a GE measurement process $p(y|\phi)$ having a trainable nonlinearity and a heteroskedastic skewed-t noise model. The resulting latent phenotype model was trained on the data of [S6]. Parameter uncertainties were estimated by repeating this inference procedure on 10 simulated datasets.

The resulting values for $\vec{\theta}_{\rm F}$ and $\vec{\theta}_{\rm B}$ are illustrated as heatmaps in Fig. 6b of the main text, along with the best estimates and standard errors for $\theta_F^0$ and $\theta_B^0$. These values are expressed in the "wild type" gauge, i.e., $\theta_F^0$ and $\theta_B^0$ are the folding and binding energies of the wild-type GB1 sequence, while the elements of the vectors $\vec{\theta}_{\rm F}$ and $\vec{\theta}_{\rm B}$ describe the energetic effects of single amino acid mutations. All of these parameter values represent Gibbs free energies computed using $k_B T = 0.582\ \frac{\rm kcal}{\rm mol}$, since the RNA display experiment of [S6] was performed at room temperature.

#### S3.3 Thermodynamic model for the *lac* promoter

Fig. S4b illustrates a thermodynamic model for transcriptional activation at the *lac* promoter of *Escherichia coli*. This model describes the DNA binding energies of two transcription factors, the cAMP receptor protein (CRP) and $\sigma^{70}$ RNA polymerase (RNAP), as well as a cooperative interaction between these two proteins that occurs when both are bound to promoter DNA. This model was proposed by [S10] and trained on data from a sort-seq MPRA in which a 75 bp region spanning the CRP and RNAP binding sites of the wild-type *lac* promoter was mutagenized at 12% per nucleotide, resulting in $\sim$9 mutations per sequence on average.

Our model assumes that promoter DNA can be in one of four possible states: (1) empty, (2) bound by CRP, (3) bound by RNAP, and (4) bound by both CRP and RNAP. State (1) is taken to be the reference state. $\Delta G_{\rm C}$ denotes the Gibbs free energy of CRP binding to DNA, and $\Delta G_{\rm R}$ is the energy of RNAP binding to DNA. Both of these quantities are assumed to be additive functions of the lac promoter sequence $x$. More specifically, $\Delta G_{\rm C}$ is assumed to depend only on a 26 nt subsequence (the CRP binding site), the one-hot encoding of which we denote $\vec{x}_{\rm C}$. Similarly, $\Delta G_{\rm R}$ is taken to depend on a 41 nt subsequence $\vec{x}_R$ representing the RNAP binding site. The corresponding parameters of these additive models are $\theta_{\rm C}^0, \vec{\theta}_{\rm C}$ for CRP, and $\theta_{\rm R}^0, \vec{\theta}_{\rm R}$ for RNAP. $\Delta G_{\rm I}$ is the Gibbs free energy of interaction between DNA-bound CRP and DNA-bound RNAP, and is assumed to be independent of sequence. The latent phenotype of interest, $\phi$, is the fraction of time that RNAP is bound to promoter DNA, a quantity that we assume is proportional to the rate of transcript initiation and thus the level of gene expression. The state-specific activities are therefore fixed and given by $\phi_{(1)} = \phi_{(2)} = 0$ and $\phi_{(3)} = \phi_{(4)} = 1$. The resulting set of trainable G-P map parameters is

$$\theta = \left\{ \theta_{\rm C}^0, \vec{\theta}_{\rm C}, \theta_{\rm R}^0, \vec{\theta}_{\rm R}, \Delta G_{\rm I} \right\}. \tag{S8}$$

This biophysical G-P map, $\phi(x;\theta)$, was paired with an MPA measurement process and trained on the data of [S10]. The resulting inferred values for $\vec{\theta}_{\rm C}$ and $\vec{\theta}_{\rm R}$ are illustrated in Fig. 6d as sequence logos; the inferred value of $\Delta G_{\rm I}$ is also shown. The standard errors on $\Delta G_{\rm I}$ were estimated by repeating the inference procedure on 10 simulated datasets.

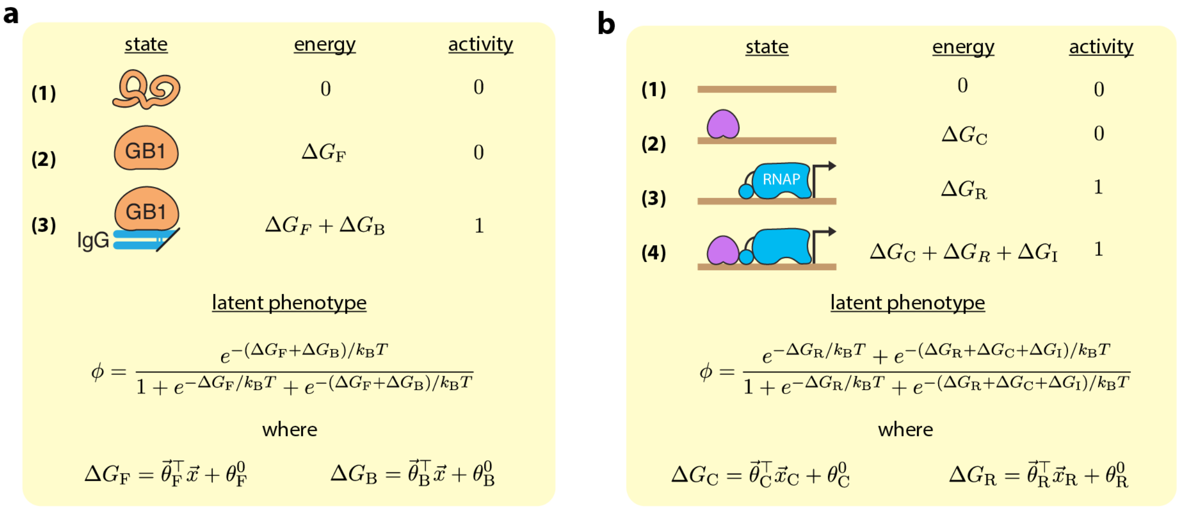


Figure S4: Thermodynamic models featured in Fig. 6, main text. **(a)** Three-state model for GB1 folding and binding to IgG. $\vec{x}$ denotes a one-hot encoding of a variant GB1 protein sequence. The trainable G-P map parameters are $\vec{\theta}_{\rm F}$, $\theta_{\rm F}^0$, $\vec{\theta}_{\rm B}$, and $\theta_{\rm B}^0$. **(b)** Four-state model for transcriptional regulation of the *E. coli lac* promoter by CRP (purple) and RNAP (blue). Variable promoter DNA is indicated in brown. $\vec{x}_C$ and $\vec{x}_R$ respectively denote one-hot encodings of a variant 26 nt CRP binding site and a variant 41 nt RNAP binding site. The trainable G-P map parameters are $\vec{\theta}_{\rm C}$, $\theta_{\rm C}^0$, $\vec{\theta}_{\rm R}$, $\theta_{\rm R}^0$, and $\Delta G_{\rm I}$.

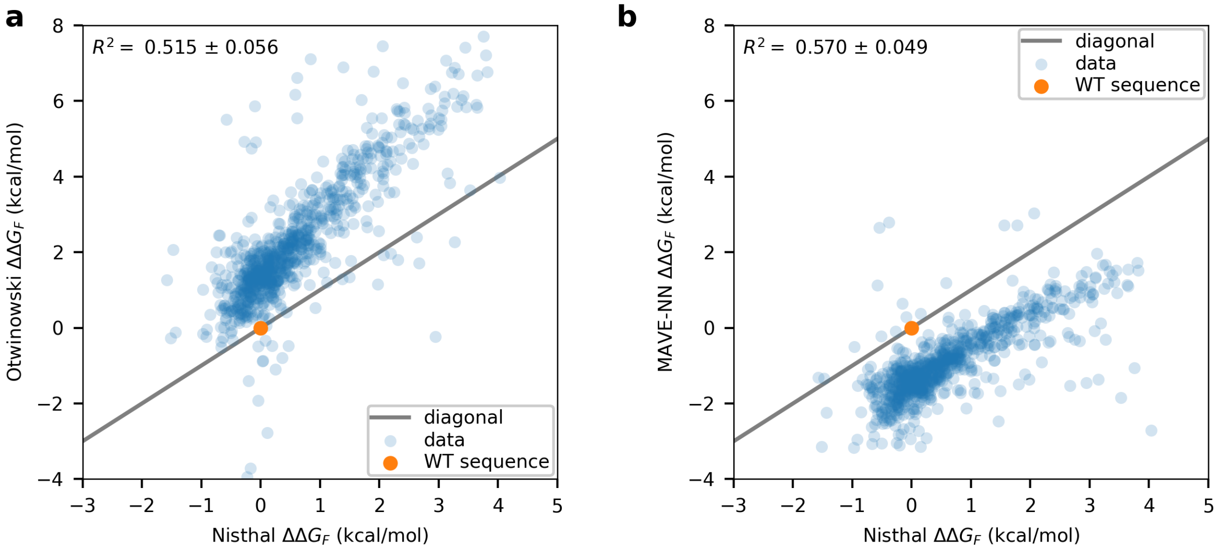


Figure S5: Gibbs free energies of folding ($\Delta G_{\rm F}$) measured by Nisthal et al. [S11] compared to predictions made by the thermodynamic models inferred by **(a)** Otwinowski [S9] and **(b)** MAVE-NN. Dotted line indicates equality between the $x$- and $y$-coordinates.

### References


[S1] D. Barber and F. Agakov, "The IM Algorithm: A variational approach to Information Maximization," *Advances in neural information processing systems.*, 2003.

[S2] J. B. Kinney and G. S. Atwal, "Parametric Inference in the Large Data Limit Using Maximally Informative Models," *Neural computation*, vol. 26, no. 4, pp. 637–653, 2014.

[S3] G. S. Atwal and J. B. Kinney, "Learning Quantitative Sequence-Function Relationships from Massively Parallel Experiments," *Journal of Statistical Physics*, vol. 162, no. 5, pp. 1203–1243, 2016.

[S4] M. Chalk, O. Marre, and G. Tkacik, "Relevant sparse codes with variational information bottleneck," *arXiv*, 2016.

[S5] A. A. Alemi, I. Fischer, J. V. Dillon, and K. Murphy, "Deep Variational Information Bottleneck," *arXiv*, 2016.

[S6] C. A. Olson, N. C. Wu, and R. Sun, "A comprehensive biophysical description of pairwise epistasis throughout an entire protein domain.," *Current biology*, vol. 24, pp. 2643 – 2651, 11 2014.

[S7] M. S. Wong, J. B. Kinney, and A. R. Krainer, "Quantitative Activity Profile and Context Dependence of All Human 5' Splice Sites.," *Molecular cell*, vol. 71, pp. 1012–1026.e3, 08 2018.

[S8] "MAVE-NN online documenataion." `https://mavenn.readthedocs.io/`.

[S9] J. Otwinowski, "Biophysical Inference of Epistasis and the Effects of Mutations on Protein Stability and Function.," *Mol Biol Evol*, vol. 35, pp. 2345 – 2354, 10 2018.

[S10] J. B. Kinney, A. Murugan, C. G. Callan, and E. C. Cox, "Using deep sequencing to characterize the biophysical mechanism of a transcriptional regulatory sequence," *Proceedings of the National Academy of Sciences*, vol. 107, pp. 9158–9163, 05 2010.

[S11] A. Nisthal, C. Y. Wang, M. L. Ary, and S. L. Mayo, "Protein stability engineering insights revealed by domain-wide comprehensive mutagenesis," *Proceedings of the National Academy of Sciences*, vol. 116, no. 33, pp. 16367–16377, 2019.
